# Characterizing the effect of short wavelengths on the floral flavonoid metabolome of medicinal cannabis using a comparative computational metabolomics workflow

**DOI:** 10.64898/2026.04.28.721290

**Authors:** Willy Contreras-Avilés, Laura Rosina Torres-Ortega, Ep Heuvelink, Leo F. M. Marcelis, Justin J. J. van der Hooft, Iris F. Kappers

**Author notes:** These authors contributed equally to this work (Willy Contreras-Avilés, Laura Rosina Torres-Ortega).

## Abstract

**Background:** Controlled-environment cultivation of medicinal cannabis (*Cannabis sativa* L.) typically optimizes light conditions to enhance the biosynthesis of pharmaceutically important metabolites like cannabinoids. Such experimental strategies may also influence other specialized metabolites like terpenoids, flavonoids, alkaloids, among others. Previous untargeted metabolomics studies testing short wavelength conditions like UV and blue light have shown that terpenoids and prenylated flavonoids in cannabis leaves respond differentially. However, since metabolomic studies in cannabis have so far mostly focused on floral cannabinoids, a comprehensive untargeted study into cannabis’ floral metabolome response to short wavelengths is currently lacking.

**Objectives:** Our study investigates the impact of short wavelength usage on cannabis specialized metabolism, and in particular the influence of UVB, UVA, and blue light on the cannabis floral flavonoid metabolome and associated glycosylation moieties.

**Methods:** Cannabis plants were grown under a white background light and exposed to supplemental UVB, UVA, or blue light during the generative phase of the cultivation cycle. Treatments were compared to a reference white background light without UV or blue light. Metabolites from floral tissue were extracted and analyzed via ultra-performance liquid chromatography-tandem mass spectrometry. A comparative metabolomics workflow was designed and used to characterize the floral flavonoid metabolome and associated glycosylation moieties.

**Results:** Our results demonstrate how short wavelengths differentially affect the metabolism of natural product compound classes including polyketides and phenylpropanoids/shikimates. Blue light induced flavonoids similarly to how UVB did, while both UVA and blue light specifically induced flavanones accumulation. UVB showed the strongest regulatory effect on flavonoids production and glycosylation patterns.

**Conclusions:** UVB reshapes the cannabis floral flavonoid metabolome by selectively stimulating the accumulation and structural modification of flavonoids. Therefore, UVB application in cannabis cultivation represents a useful horticultural strategy to increase inflorescence medicinal quality without affecting cannabinoid levels.

## Introduction

*Cannabis sativa* L. is a medicinal plant producing health-affecting specialized metabolites including cannabinoids, terpenoids, flavonoids, as well as unique flavol-alkaloids and prenylated flavones (cannflavins) (Jin et al., 2020; Muller & De Villiers, 2025; Rea et al., 2019). Cannabinoids and terpenoids predominantly accumulate in cannabis inflorescences, whereas flavonoids, flavo-alkaloids, and cannflavins also can be found in leaves (Jin et al., 2020). The accumulation of cannabis specialized metabolites can be influenced in controlled-environment agricultural systems where light and other environmental factors are largely under control (Contreras-Avilés et al., 2024; Desaulniers Brousseau et al., 2021). Light is an essential environmental factor consisting of photons that once perceived by cryptochromes (CRYs) and phytochromes (PHYs) trigger a signalling cascade driving photomorphogenesis and specialized metabolism (McCree, 1981; Opálková et al., 2018). Shorter wavelengths including UVB (280 - 315 nm) and UVA (315 - 400 nm) influence growth, morphology, and plant specialized metabolism via UV RESISTANCE LOCUS 8 (UVR8) and CRYs photoreceptors, respectively (Supplementary Notes 1) (Contreras-Avilés et al., 2024; Rai et al., 2021; Sun et al., 2024, 2025). Specialized metabolites are of great anthropocentric interest due to their nutritional and medicinal properties (Supplementary Notes 2) (Contreras-Avilés et al., 2024). The flavonoids specialized metabolite class (Supplementary Notes 3) is widely induced under UV radiation (Rai et al., 2021), and has been described to have multifunctional roles in plants including photoprotection, antioxidant activity, and signalling (Agati & Tattini, 2010; Mao et al., 2025; Patil et al., 2024). Flavonoids originate from the phenylpropanoid pathway, where phenylalanine-derived *p*-coumaroyl-CoA is combined with malonyl-CoA units via chalcone synthase to form the core flavonoid skeleton. Subsequent enzymatic modifications diversify this core structure into the major flavonoid subclasses flavanones, flavanols, anthocyanins, flavonols, and (iso)flavones, which are often found as hydrophilic glycosides or lipophilic methyl/prenylated/geranylated molecules (Shomali et al., 2022). Among flavonoids, flavonols function as specialized metabolites that mitigate short-wavelengths-induced stress through reactive oxygen species (ROS) scavenging, metal chelation, inhibiting ROS-generating enzymes, and as cofactors of antioxidant enzymes (Shomali et al., 2022). UVB has been shown to induce flavonols as kaempferol in *Coriandrum sativum* (Fraser et al., 2017) and quercetin in *Ginkgo biloba* (Zhao et al., 2020) as well as glycosylated derivatives (Supplementary Notes 4) *Asparagus officinalis* L. (Eichholz et al., 2012). Similarly, increased solar radiation enhanced quercetin-3-O-rutinoside and luteolin-7-O-glucoside in *Ligustrum vulgare* (Tattini et al., 2004), while combined UV (UVB + UVA) and blue light promoted flavonol disaccharides in *Cucumis sativus* (Palma et al., 2021).

Short wavelengths (<500 nm) promote ROS and regulate accumulation of glycosylated flavonoids via UVR8 signalling (Markovitsi et al., 2013; Nishigori et al., 2023; Rai et al., 2021). However, responses are species and structure-dependent, for example, UV increased kaempferol glycosides but decreased quercetin glycosides in *Brassica oleracea* (Moreira-Rodríguez et al., 2017; Rechner et al., 2016), while shifting mono- and- di-glycosylated flavonoids in *Glycyrrhiza uralensis* (Zhang et al., 2018). UVB enhanced either kaempferol glycosides or quercetin glycosides depending on species (*Brassica rapa, B. oleracea, B. carinata*) (Neugart & Bumke-Vogt, 2021). UVA increased kaempferol glycosides in *Glycine max*, (Lim et al., 2021), and flavone glycosides in *Scutellaria baicalensis* (Miao et al., 2022), while blue light promoted flavones glycosides in cereals and anthocyanins in *Malus domestica* (Kokalj et al., 2019; Muthusamy et al., 2020). Overall, glycosylated flavonoid responses appear structure-dependent (Neugart & Schreiner, 2018), and it remains unclear whether UVA and blue light elicit distinct effects despite sharing CRYs signalling pathways (Rai et al., 2019).

Few studies have investigated the effects of UV and/or blue light on flavonoid accumulation in cannabis, and those available lack detailed analysis of flavonoid profiles and glycosylation patterns (Kotiranta et al., 2024; Marti et al., 2014). Although flavonoids are abundant in leaves, their presence in inflorescences is cultivar-dependent or associated with oxidative stress (Desaulniers Brousseau et al., 2021), and their physiological role in light responses remains unclear. Computational metabolomics (“cannabinomics”) has been used to characterize cannabis chemical diversity supporting chemovar classification, breeding, and bioactivity studies (Aliferis & Bernard-Perron, 2020; Cerrato et al., 2023; Liu et al., 2025; Myoli et al., 2024a, 2024b; Tang et al., 2023; Vásquez-Ocmín et al., 2021), but these approaches primarily focus on cannabinoids and terpenoids, often overlooking flavonoids. Untargeted UPLC–MS/MS can yield thousands of chromatographic features for comparative analyses, but translating these signals into confident chemical identities remains a key bottleneck, particularly for flavonoids and their glycosylated forms, often occurring as positional isomers with highly similar MS/MS patterns and limited spectral library coverage (Bittremieux et al., 2022; Li et al., 2024). Therefore, using cannabis inflorescences from a prior short-wavelength exposure experiment, we created and applied an integrated computational workflow (mzmine–MN–MS2LDA–SIRIUS–MS2Query–DreaMS) to characterize the floral flavonoid metabolome, linking differential abundance patterns to molecular families (MN), substructures and glycosylation moieties (MS2LDA), chemical class assignments (SIRIUS/CANOPUS), and structural analogues via machine and deep learning similarity (MS2Query; DreaMS) (Supplementary Notes 5) (Bushuiev et al., 2025; De Jonge et al., 2023; Dührkop et al., 2019; Torres Ortega et al., 2025). We hypothesize that short wavelengths promote flavonoid glycosylation as an antioxidant strategy under oxidative stress and our results specifically show that UVB induces flavonol accumulation and enhances glycosylation, while UVA and blue light elicit subclass-specific responses.

## Materials and methods

We used cannabis inflorescences collected in a previous study (Contreras-Aviles et al., submitted). In short: plants were grown for 12 days under a white (10% Blue, 10% Green, 80% Red) background light and a 18 hrs photoperiod (400 µmol m^-2^ s^-1^) to stimulate vegetative growth, after which the flowering phase was initiated by switching to a 12 hrs photoperiod (600 µmol m^-2^ s^-1^). Light treatments started 14 days after the short-day phase was initiated by exposing plants to supplemental UVB (2 µmol m^-2^ s^-1^; 3 hrs d^-1^), UVA (88 µmol m^-2^ s^-1^; 12 hrs d^-1^), or blue (84 µmol m^-2^ s^-1^; 12 hrs d^-1^) light; all treatments were applied at the beginning of the photoperiod. UVA and blue intensity and photoperiod were matched to ensure comparability, while UVB parameters were selected to minimize harm while stimulating specialized metabolism. All treatments were compared to a reference without supplemental light. To evaluate the coloration of glandular trichomes, these anatomical structures were imaged (Supplementary Information 6).

### Ultra-high-performance liquid chromatography mass spectrometry (UHPLC-MS/MS) based metabolomic analysis

At maturity (eight weeks after the onset of the generative phase) inflorescence samples were taken from within 5 cm of the apical section of the top inflorescences and flash frozen in liquid nitrogen. Frozen samples were freeze-dried (Alpha 1-4 LSCbasic freeze-dryer, Martin Christ, Osterode am Harz, Germany) for 36 hours and ground into a fine powder (MM 200 mixer mill, Retsch, Dale I Sunnfjord, Norway). After sonication (Branson 2800 ultrasonic bath), combined supernatants were dried (SpeedVac SPD2030-230, Savant, Thermo Scientific, USA) and reconstituted in 80%MeOH(0.01% formic acid); sonicated for 30 seconds and centrifuged at 21,000×g at 4°C for 10 min prior to injection (Supplementary Information 7).

### Data preprocessing and metabolite feature finding

The acquired raw datasets (raw, ESI positive and negative ionization mode centroid data) were converted to .mzML using ProteoWizard MSConvert (3.0) and pre-processed using mzmine 4.7.8. (Supplementary Information 8). After filtering, 5,680 (positive ionization mode) and 8,664 (negative ionization mode) features were retained.

### Metabolite and substructure *in silico* annotation

The output from mzmine was used for *in silico* annotation using SIRIUS (Dührkop et al., 2019) for molecular formulas and ZODIAC (8.5.6) (Ludwig et al., 2020) to refine assignments. Chemical class annotations were assigned with CANOPUS as a part of the SIRIUS framework (Dührkop et al., 2019). The settings used for SIRIUS are described in Supplementary Information 7. To complement these annotations, we applied two similarity-based spectral annotation approaches. First, MS2Query (De Jonge et al., 2023) was used to retrieve candidate library annotations for each query spectrum based on learned MS/MS similarity and re-ranking, providing putative identifications. Second, DreaMS (Bushuiev et al., 2025) was applied as an embedding model to perform nearest-neighbour spectra similarity, obtaining the most similar reference spectra above a cosine similarity threshold of 0.75 (via the DreaMS web interface). The output of both tools includes the chemical classes inferred by applying NPClassifier to the associated candidate structures and nearest neighbour annotations (Kim et al., 2021) (Figure 1). Substructure-level annotation was done using MS2LDA 2.0 (Torres Ortega et al., 2025) (more details Supplementary Information 9).

**Figure 1.**
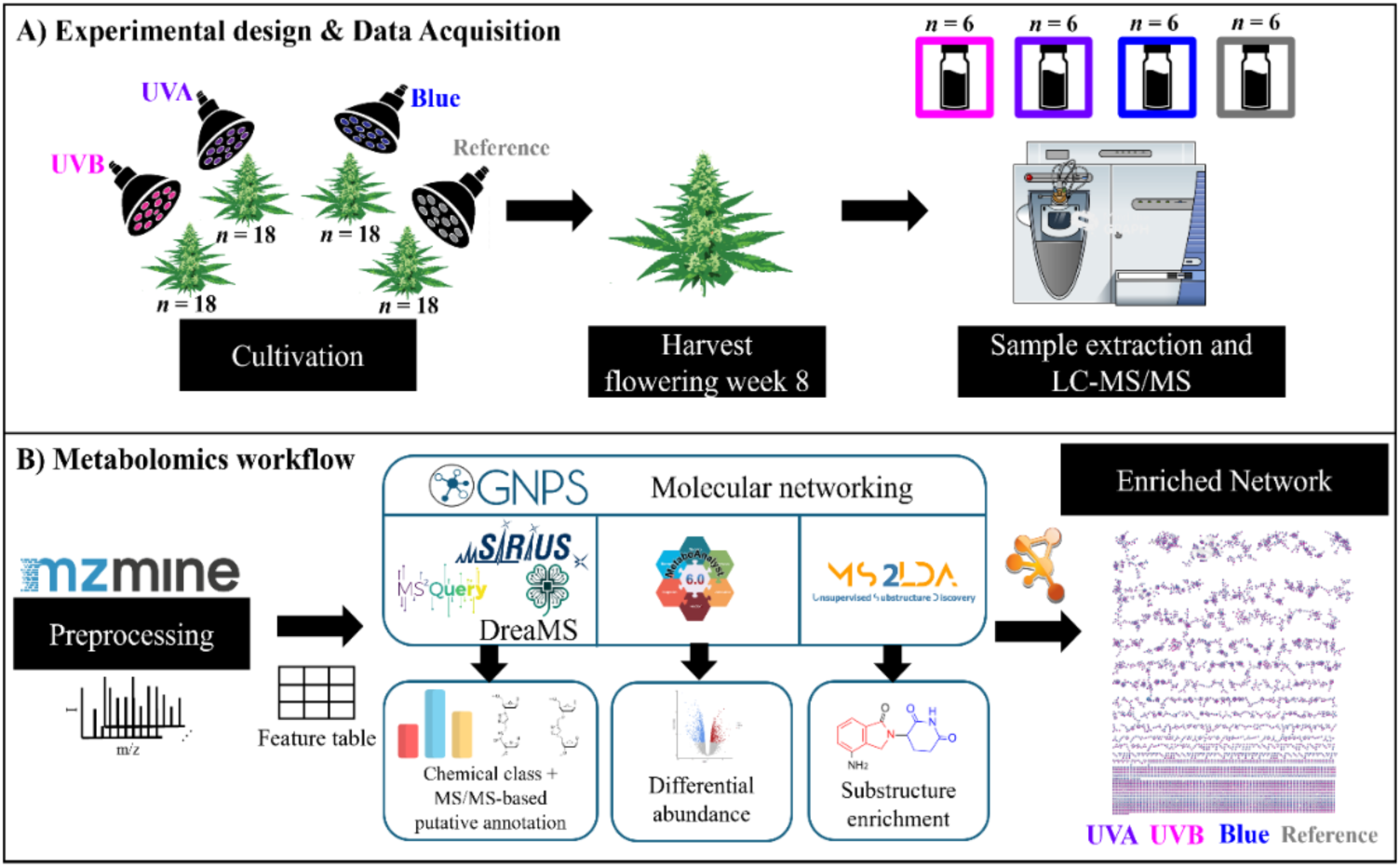
Comparative computational metabolomic workflow. Building on a previous short-wavelength experiment, A) cannabis plants were exposed to UVB, UVA, and blue light treatments, and inflorescence samples were taken eight weeks after the onset of flowering, followed by UPLC-MS/MS analysis. B) mzmine was used to preprocess and produce a feature table which was also complementarily used for GNPS FBMN and MS/MS-based putative annotation (SIRIUS/CANOPUS. MS2Query, DreaMS). Differential abundance analysis (MetaboAnalyst) and substructure discovery (MS2LDA) were also performed. Altogether, the results were integrated into an enriched network visualized in Cytoscape prioritizing flavonoid metabolites features.

### Feature-base molecular networking

Feature-based molecular networking (FBMN) was performed in GNPS2 (Nothias et al., 2020) using the mzmine .mgf, the feature quantification table, and a sample metadata file as inputs. The settings for the spectral library matching are described in Supplementary Notes 10. The resulting network was exported as a GraphML file and imported into Cytoscape (v3.10.3) (Shannon et al., 2003) for visualization, annotation outputs were integrated as node attribute tables in Cytoscape using the shared feature identifier (Supplementary Information 10).

### Chemical classification and chemometric analysis

The experiment was set up and analysed as complete randomized design, where six biological samples were considered for every treatment. Five dataset tables were generated from positive ionization data including: *global* (all features), *phenylpropanoid/shikimate*, *phenylpropanoid*, *flavonoids*, and *final annotation* (manually annotated flavones, flavonols, and anthocyanins, with duplicates and isomers removed). Datasets were pre-processed in MetaboAnalyst 13.0 (Pang et al., 2021) using left-censored data estimation, median normalization, log₁₀ transformation, and Pareto scaling. Partial least squares discriminant analysis (PLS-DA) with five-fold cross-validation, variable in projection (VIP) scores, and hierarchical clustering heatmaps were used to visualize treatment effects. Features with VIP ≥ 1.5 were considered significant, and ≥ 2.5 as strongest drivers. Mean separation of relative abundances was performed by one-way ANOVA with Fisher’s protected LSD test (α = 0.05), and treatment effect magnitude was assessed by volcano plot analysis (more details in Supplementary Information 11).

## Results

A comparative metabolomic workflow was applied to characterize the cannabis floral metabolome. We first explored the mapped treatment-responsive chemical space using feature-based molecular networking and class-level annotations to identify molecules most affected by UVB, UVA, and blue light. Secondly, we focused on the phenylpropanoid/shikimate pathway to identify various chemical compound classes connected to this pathway and compared their relative abundances in response to the different short-wavelength treatments. Thirdly, we resolved flavonoid subclasses and glycosylation types (O- vs C-glycosides) by combining structural information obtained from molecular family grouping, substructure motifs, and candidate structure annotations.

### Metabolomic profiling of cannabis floral metabolome

The spectra were preprocessed using mzmine to obtain the quantitative abundance of the metabolic features, and the global feature-based molecular network generated from the positive-ionisation data contained 5,700 features connected by 7,400 edges (Figure 2A) revealed multiple molecular families covering diverse biosynthetic pathways. Based on VIP scores ≥ 1.5, short-wavelength treatments affect the cannabis metabolome in seven major chemical classes: polyketides, amino acids and peptides, terpenoids, fatty acids, alkaloids, carbohydrates, and phenylpropanoid/shikimates, according to NPClassifier annotation (Figure 2B). Across the network, 42% of nodes had a match to entries in public structural and spectral libraries, indicating a substantial uncharacterised chemical space in cannabis flowers.

**Figure 2.**
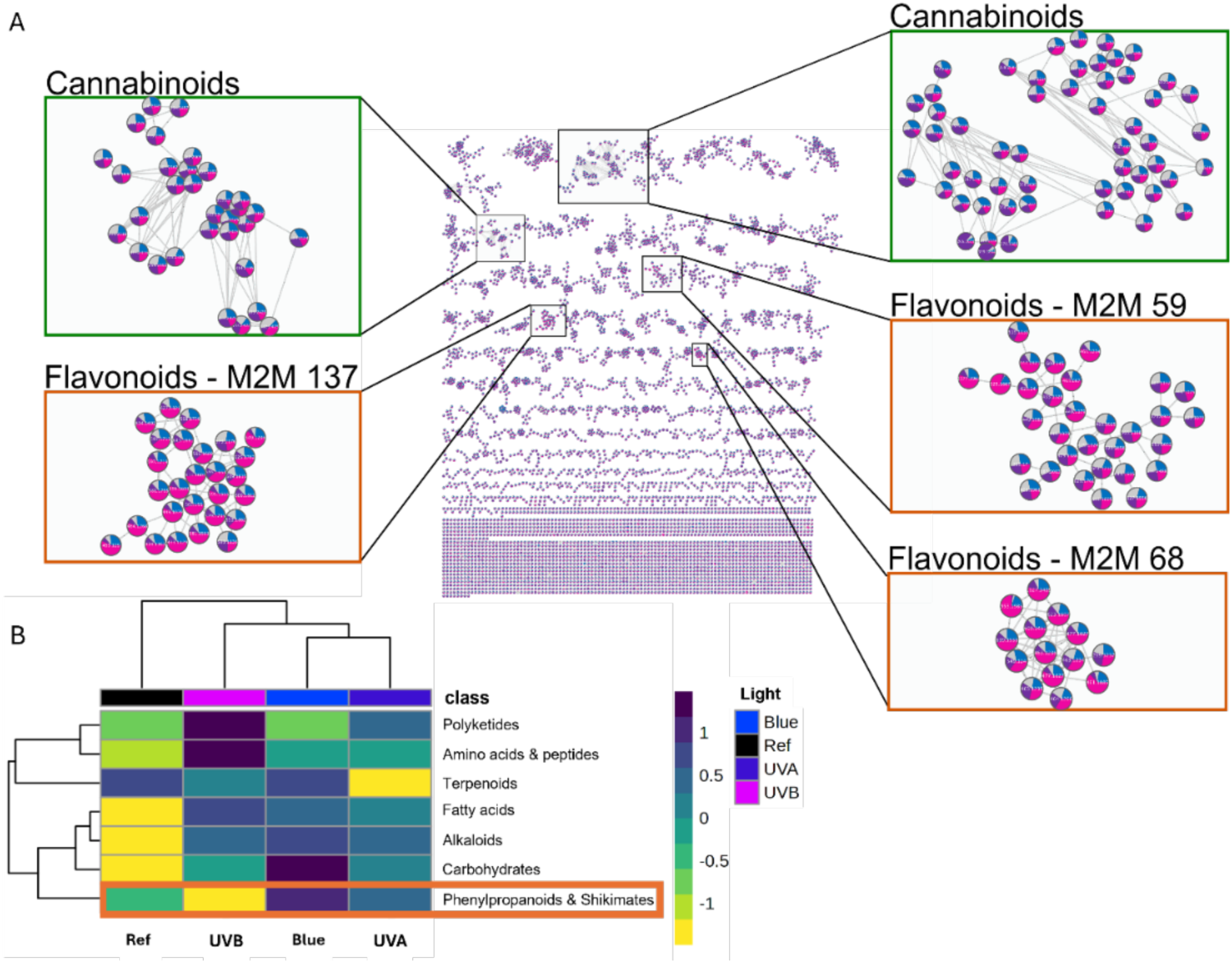
Effect of short wavelengths (UVB, UVA, blue light) on the global metabolome in *Cannabis sativa* L. inflorescences collected eight weeks after the onset of flowering. (A) Feature-based molecular network (FBMN) in positive ionisation mode: mass spectral molecular network of the flowers indicating flavonoids and cannabinoids clusters. Pie charts represent relative abundances for each feature (compound) according to the feature-based molecular network in positive ionization (FBMN-POS), where each color represents UVB (pink), UVA (purple), blue (blue), and reference (grey) short-wavelength treatments. (B) Heatmap based on the hierarchical clustering of the ‘*global*’ dataset of the floral metabolome with variables in projection (VIP) score ≥ 1.5; the yellow to dark blue bar corresponds to a Z-scaling ranging from -1 to 1; the orange rectangle highlights the phenylpropanoid/shikimate subgroup.

To increase putative coverage beyond library matches, we applied complementary *in silico* tools. SIRIUS returned 183 features (3.2%) with structures above 0.6 of confidence score, a commonly used confidence threshold for formula-to-structure assignments in datasets with limited spectral coverage (Dührkop et al., 2019). MS2Query provided analogue-level similarity (score > 0.8) for 22.35% of features. Using DreaMS (also for analogues) with a similarity cut-off of 0.75, annotations for 264 features (4.61%) were obtained. Together, these approaches expanded the interpretable cannabis inflorescence chemical space while still reflecting that a large fraction of the floral metabolome remains only partially characterised.

The negative-ion FBMN contained 8,664 nodes, but only 3.5% of these had a library match. In addition, MS2LDA Mass2Motif Annotation Guidance (MAG) was not available for negative ionization mode at the time of analysis. We therefore focused on the positive-ion network for subsequent analysis.

Given their medicinal relevance, we first evaluated cannabinoid-related molecular families. Cannabinoid features did not differentiate across short-wavelength treatments (Figure 2A), indicating they did not respond in this study. Cannabinoid substructures were nonetheless captured by several Mass2Motifs: in molecular family MF1, motif_102 and motif_120 associated with features annotated as cannabidiol (CBD) and cannabigerol (CBG), both containing the fragment 193.12 m/z from side-chain cleavage of the six-carbon ring (Citti et al., 2019; Ferrer, 2020) (Supplementary Figure 1 A,B). In MF2, motif_44 and motif_9 associated with cannabidiolic (CBDA) and cannabigerolic acid (CBGA) features (Supplementary Figure 1C,D), where motif_44 is dominated by the 341.21 m/z fragment (water loss from the carboxylic acid) and motif_9 by 219.10 m/z (C–C bond cleavage)

Given our central hypothesis regarding short-wavelength regulation of flavonoid-like molecules, we next prioritised phenylpropanoid/shikimate-related molecular families that were both treatment-responsive (VIP ≥ 1.5) and enriched in flavonoid-like substructures. Substructure analysis with MS2LDA identified 150 Mass2Motifs across the dataset. Notably, one of the most frequent motifs (motif_87) (Supplementary Figure 2) returned a recommended structure corresponding to four linked glycosidic moieties, with motif containing fragments related to a hexose moiety and present in 849 spectra. From MotifDB (database with manual annotations) we obtained a hit to “fragments indicative of a glycosylation” for motif_24 and motif_45 from MassBank and GNPS respectively with a score of 0.396. As this motif occurred broadly without a flavonoid core, it could imply one or possibly more sugar conjugations across multiple pathways in floral specialised metabolism.

The dominant Mass2Motifs enriched among 42 flavonoid-related nodes and particularly informative for resolving phenylpropanoid/shikimate responses include motif_59, motif_137, motif_68, and motif_105 (Supplementary Figure 3-6). Motif_59 captured the canonical flavonoid scaffold and occurred in aglycones such as luteolin and kaempferol (diagnostic fragment around *m/z* 287), matching MassBank motif_42 manually annotated as “fragments indicative for kaempferol” (score of 0.5875), strengthening our assignment. Motif_137 associates with glycosylated flavonoids, showing diagnostic sugar-related fragments (e.g., *m/z* 329), and was repeatedly detected in UVB-responsive nodes with low-score MotifDB hits to MassBank motif_16 (’quercetin/glycosylated quercetin fragments’, 0.28) and motif_42 (’kaempferol fragments’, 0.28)”. Motif_68 reflected a flavonoid-like ring system with a characteristic fragment around *m/z* 301. Here, MotifDB resulted in a low score hit annotated as “5-Hydroxy-2 2-dimethyl-4-oxo-3 4-dihydro-2H-chromen-7-yl”. Finally, motif_105 represented simplified aromatic-ring substructures (*m/z* 127, 135, 139), and also matched MassBank motif_42 (’kaempferol fragments’, 0.23).

Besides flavonoid-related Mass2Motifs, we have checked the motifs with the highest degree of associated number of spectra (apart from motif_87) across the entire dataset: motif_139, motif_114, motif_115, and motif_15 (Supplementary Figure 7A-D). Motif_139 (in 1,330 spectra) was consistent with a six-membered ring containing an alkyl side chain with multiple optimised fragments, when comparing to MotifDB, the top match was annotated as “diterpenoid” (score = 0.277) from the Euphorbia Plant Mass2 MotifSet. Motif_114 was detected in 585 spectra, containing two highest losses 18.01 and 36.02, with low probabilities, 0.35 and 0.34 respectively. With a lack optimised fragments or loses, we propose this may represent a noisy Mass2Motif, as we are unable to associate to a substructure. Motif_115 represented a glycosylation by MAG, which correlated with the hits against MotifDB as for motif_87 (motif_45 and motif_24), indicative for glycosylation with a score of 0.574. Finally, for motif_15 (433 spectra) the three MAG recommendations contain a *p*-coumaroyl ester typically producing the diagnostic fragments 147.04 *m/z* and with the loss of CO the 119.04 *m/z* fragment; from MotifDB the hits were motif_20 and motif_37 from GNPS and MassBank (same annotation) described as “fragments for cinnamic/hydroxycinnamic acid substructures”. As *p*-coumaroyl ester is a hydroxycinnamic acid conjugated with an ester, this motif is therefore well annotated by MS2LDA. Altogether, we were able to see the effect of glycosylation in the most frequent motifs found across the dataset (Supplementary Table 1).

### Flavonoid abundances are differentially affected by UVB, UVA, and blue light

After identifying enriched flavonoid motifs, we assessed the effect of short wavelengths on the accumulation of distinct flavonoid subclasses. Notably, UVB exposure was associated with a magenta coloration in glandular trichomes on bracts (Figure 3A), a phenotypic change absent under the other treatments (Supplementary Figure 8A-C). PLS-DA of the phenylpropanoid/shikimate dataset generated a model in which the first two components significantly contributed to clustering based on short-wavelength treatment (Component 1: 7.8%; Component 2: 16.5%) (Figure 3B). Model quality metrics confirmed the robust performance R^2^ = 0.98, Q^2^ = 0.52, and accuracy 0.84 (Supplementary Figure 9A). A total of 15 features with VIP scores ≥ 1.5 contributed to separation of clusters observed in the PLS-DA (Supplementary Figure 9B). From those metabolite features, six belong to the flavonoid subclass, including two flavones (ID: 2523; 2291 | VIP: 3.4, 2.8 | mass: 303.049 *m/z*; 595.165 *m/z*), two flavonols (ID: 2262; 2617 | VIP: 4.6; 3.3 | mass: 231.049 *m/z*; 463.087 *m/z*), one anthocyanidin (ID: 2627; VIP: 2.9; mass: 1177.396 *m/z*), and one flavanone (ID: 5377; VIP: 2.5; mass: 371.149 *m/z*) (Figure 3D). A clustering heat map of the phenylpropanoid dataset revealed that the flavonoids subclass showed similar abundances in response to UVB and blue light (Figure 3C), which is in accordance with previous findings on total leaf flavonoids (Kotiranta et al., 2024).

**Figure 3.**
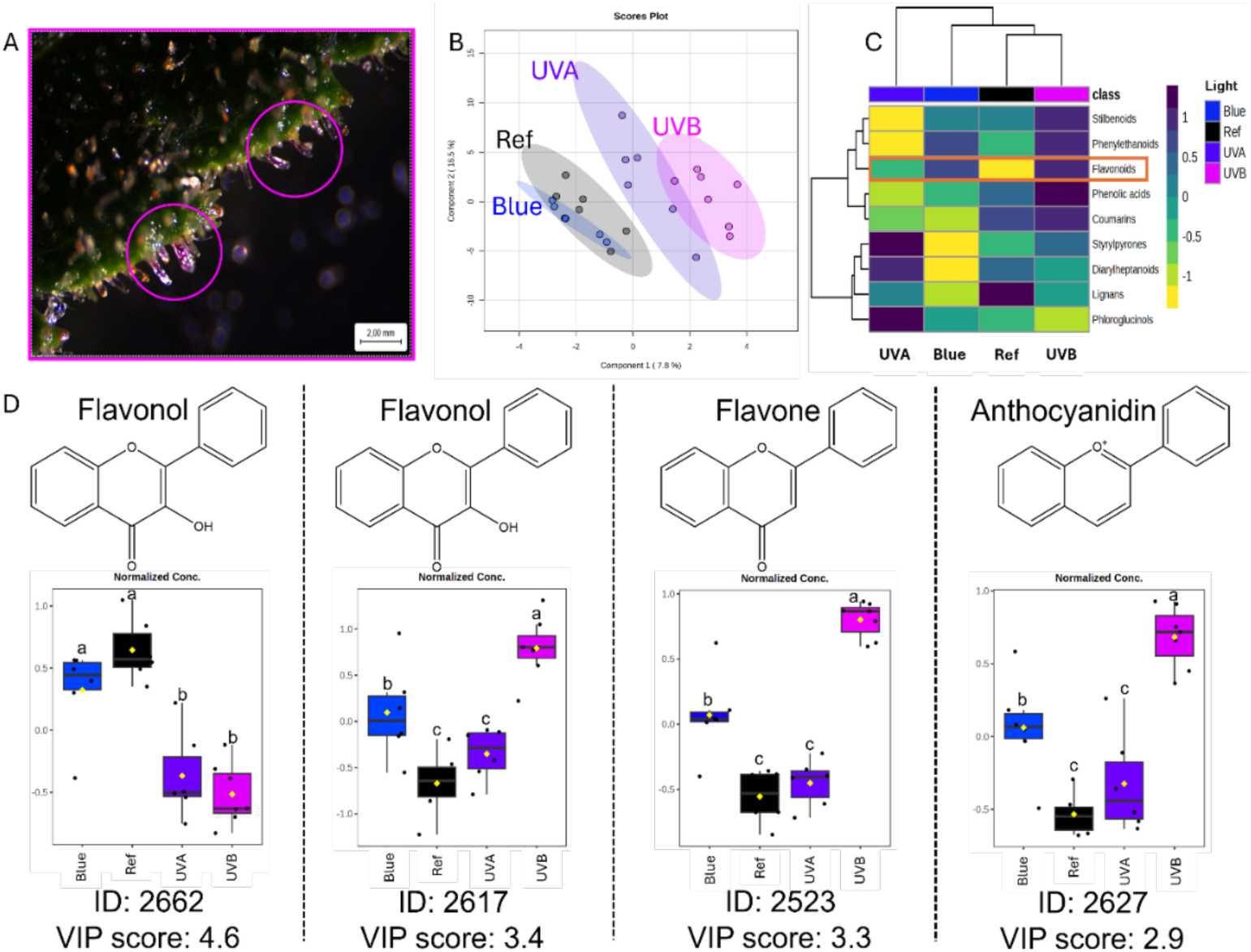
Effect of short-wavelength (UVB, UVA, blue light) on flavonoids production in *Cannabis sativa* L. inflorescences (A) Glandular trichomes on bracts in inflorescences of generative plants (eight weeks after onset of flowering) exposed to UVB. Pink circles indicate glandular trichomes with magenta coloration; (B) Partial least square - discriminant analysis (PLS-DA) of the phenylpropanoid/shikimate dataset (cross-validation R^2^ = 0.98, Q^2^ = 0.52). Ellipses represent 95% confidence region for each short-wavelength treatment; (C) Heatmap based on hierarchical clustering of the ‘*phenylpropanoid*’ dataset with variables in projection (VIP) score ≥ 0.5; yellow to dark blue bar corresponds to a Z-scaling ranging from -1 to 1, orange rectangle highlights the flavonoids subclass; (D) Relative abundances of flavonoid features within the ‘*phenylpropanoid/shikimate*’ dataset with VIP scores ≥ 2.5 in inflorescences exposed to UVB, UVA, or blue light. Positive or negative values on the x-axis indicate higher and lower relative abundance compared with the dataset’s mean. Boxplots represent median (horizontal line), interquartile range (box), mean (yellow diamond), maximum and minimum (whiskers), and outliers (beyond whiskers). Mean values represent the average of six inflorescences per treatment (n = 6). Different letters indicate significantly different values for short-wavelength treatments according to a Fisher’s protected LSD test (α = 0.05).

Comparing the different short-wavelength treatments to the reference revealed that UVB significantly increased relative abundances of four flavonoids including two flavones (ID: 2523; 6168), a flavonol (ID: 2617), and anthocyanidin (ID: 2627), whereas it also decreased the abundance of the flavonol (ID: 2662) (Table 1; Supplementary Figure 10A). UVA significantly increased relative abundances of six flavonoids including two flavones (ID: 2291; 5265) and two flavanones (ID: 5377; 5352) and decreased a flavone (ID: 2435 | Log2FC: -1.1) and flavonol (ID: 2262) (Table 1; Supplementary Figure 10B). Finally, blue light significantly increased relative abundances of four flavonoids including flavonol (ID: 2617), flavone (ID: 2523), anthocyanidin (ID: 2627), and flavanone (ID: 5377) (Table 1; Supplementary Figure 10C). Altogether, these results suggest that among identified flavonoid classes, flavones and flavonols are the most strongly induced in response to short wavelengths. UVB and blue light upregulate similar flavonoid subclasses (flavonol, flavone, anthocyanidin), whereas UVB and UVA downregulate flavonol accumulation. In contrast, UVA and blue light promote flavanone production. Overall, the magnitude of treatment-associated changes was greatest under UVB, with blue showing partial overlap and UVA producing a distinct subclass profile.

**Table 1.** Effect of short-wavelength (UVB, UVA, blue light) on flavonoids relative abundances in inflorescences from eight weeks after the onset of flowering of *Cannabis sativa* L.

| ID | VIP Score<sup>#</sup> | Class | Mass (*m/z*) | Log<sub>2</sub>FC<sup>§</sup> |  |  |
| --- | --- | --- | --- | --- | --- | --- |
|  |  |  |  | UVB vs Ref | UVA vs Ref | Blue vs Ref |
| 2662 | 4.6 | Flavonol | 625.176 | -2.1*** | -1.9*** | - |
| 2617 | 3.4 | Flavonol | 463.087 | 3.2*** | - | 2.0** |
| 2523 | 3.3 | Flavone | 303.049 | 2.7*** | - | 1.3** |
| 2627 | 2.9 | Anthocyanidin | 1177.396 | 2.3*** | - | 1.2** |
| 2291 | 2.8 | Flavone | 595.165 | - | 1.6*** | - |
| 5377 | 2.5 | Flavanone | 371.149 | - | 1.4*** | 1.2* |
| 6168 | 2.1 | Flavone | 283.062 | 1.1* | - | - |
| 5265 | 1.9 | Flavone | 469.185 | - | 2.8*** | - |
| 2435 | 1.7 | Flavone | 433.112 | - | -1.1* | - |
| 5352 | 1.5 | Flavanone | 469.185 | - | 5.8*** | - |

§ Log<sub>2</sub>fold-change (FC) values obtained from volcano plot analysis where the short-wavelength treatments were compared to the reference. Positive and negative values indicate upregulation and downregulation of relative abundances, respectively. # Variable in projection (VIP) scores obtained from the partial least squared – discriminant analysis (PLS-DA) of the phenylpropanoid/shikimate dataset. \* *p*-value < 0.05 | \*\* *p*-value < 0.01 | \*\*\* *p*-value < 0.001.

Notably, the most strongly UVB-associated features within the phenylpropanoid/shikimate class were higher-mass nodes and flavonoid-like signals, suggesting that post-biosynthetic modification (including glycosylation) may be a major driver of the UVB response. We therefore next examined flavonoid subclasses and glycosylation patterns in detail.

### Short wavelength-dependent accumulation of glycosylated flavonoids in cannabis inflorescences

To assess the effect of UVB, UVA, and blue light on the accumulation of flavonoids subclasses, we focused on identifying glycosylation patterns associated to flavanols, flavones, and anthocyanins. Several members of the flavonoid biosynthetic pathway were detected in the cannabis floral metabolome, belonging to downstream branches originating from phenylalanine (Figure 4). Some of the early intermediates in the flavonoid pathway including apigenin, eriodictyol, and dihydrokaempferol could be identified, but were not differentially affected by UV or blue light treatments. Numerous flavones and flavonols were identified, predominantly in glycosylated forms.

**Figure 4.**
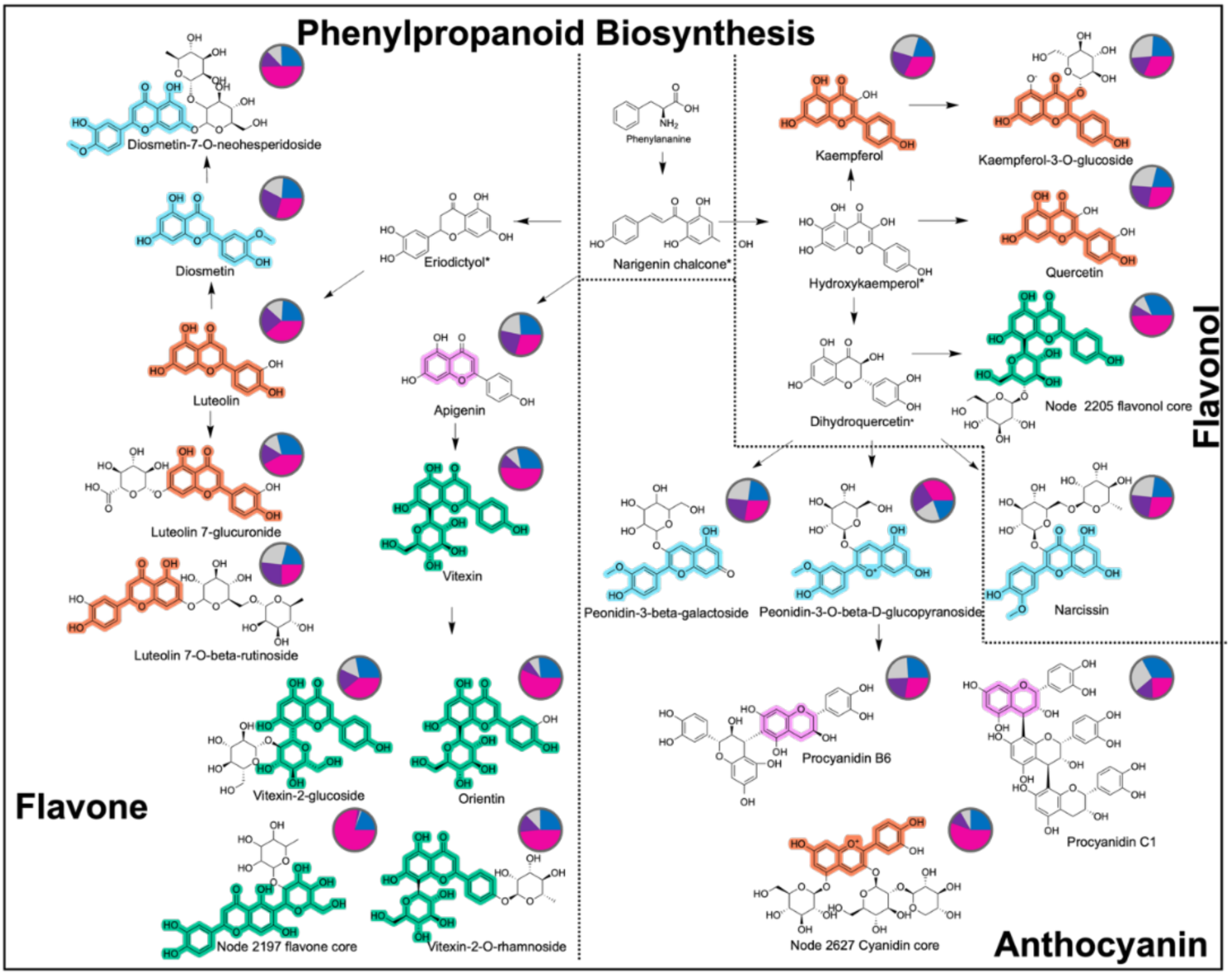
Effect of short-wavelength (UVB, UVA, blue light) on the biosynthesis of flavones, flavonols, and anthocyanins in *Cannabis sativa* L. inflorescences. A simplified overview of the phenylpropanoid biosynthesis focused in the downstream section responsible to produce flavonoids. Chemical structures highlighted in light-blue, orange, green and pink correspond to motif_68, motif_59, motif_137, and motif_105, respectively, representing shared substructures between flavonoid subclass (flavone, flavonol, and anthocyanin). Pie charts represent relative abundances for each identified feature (compound) according to the feature-based molecular network in positive ionization (FBMN-POS), where each color represent UVB (pink), UVA (purple) and blue (blue) light treatments, and reference or control (grey).

Comparative analysis of flavonoid relative abundances across short-wavelength treatments revealed a clear divergence between aglycones and glycosylated derivatives (Figure 4). Non-glycosylated flavonoids, including luteolin, kaempferol, and chrysoeriol, did not exhibit significant differences in relative abundances in response to UVB, UVA, and blue light treatments. In contrast, several glycosylated flavonoids, such as luteolin-2-O-glucoside, kaempferol-3-O-glucoside, vitexin, and orientin, showed markedly higher abundances under UVB exposure (Figure 3). These results indicate UVB preferentially promotes flavonoid glycosylation rather than increasing abundances of the corresponding aglycone backbones.

Importantly, both O-glycosylated flavonols (e.g., luteolin-2-O-glucoside and kaempferol-3-O-glucoside) and C-glycosylated flavones (e.g., vitexin and orientin) were enhanced under UVB. This concurrent accumulation of distinct glycosylation types suggests that UVB exposure broadly affects flavonoid glycosylation across structurally and biosynthetically distinct subclasses, rather than selectively inducing a single glycosylation route.

Metabolic features displaying the highest relative abundances under UVB were predominantly glycosylated and frequently of high molecular mass, most notably motif_137 (annotated as glycosylated flavonoid substructure), present in the flavonoid molecular family. Among these were orientin, luteolin-2-O-glucoside, vitexin, and kaempferol-3-O-glucoside, as well as three high-mass molecular network nodes (ID: 2205; 2197; 2627), all exceeding 1000 Da (Figure 3). Metabolic feature 2205 (precursor *m/z* 1029.344) showed a library match to O-β-D-xylopyranosylorientin and structurally related analogues, although no exact mass match was obtained. Metabolic feature 2197 (*m/z* 1189.323) lacked direct library matches but was annotated by MS2Query analogue search as a highly oxygenated flavone with multiple sugar moieties (“Flavone base + 4O, C-Hex-dHex”), and contained motif_137, supporting its classification as a glycosylated flavonoid. Metabolic feature 2627 (*m/z* 1177.396) showed a putative annotation as a highly glycosylated flavonoid or anthocyanidin-like structures and contained motif_59, reinforcing its placement within the broader flavonoid family. Although precise structural elucidation of these high-mass compounds was not achieved, their shared substructural features and strong responsiveness to UVB exposure indicate a coordinated accumulation of extensively glycosylated flavonoids.

Collectively, the combined abundance patterns and substructure-level evidence demonstrate that UVB selectively enhances both the abundance and structural complexity of glycosylated flavonoids in cannabis inflorescences. The enrichment of both C- and O-glycosylated flavonoids under UVB supports the hypothesis that UVB-driven modulation of the floral metabolome operates primarily through post-biosynthetic diversification of flavonoids, potentially expanding the antioxidant pool available for cellular compartmentalization and redox homeostasis under UV-induced stress.

## Discussion

Here, we demonstrated how a comparative computational metabolomic workflow characterized the cannabis floral metabolome by library annotations based on analogues, chemical classes predictions, and sub-structural scaffolds assignments. Based on VIP scores ≥ 1.5, short wavelength treatments affected at least seven major biochemical classes including polyketides, amino acids and peptides, terpenoids, fatty acids, alkaloids, carbohydrates, and phenylpropanoid/shikimates (Supplementary Notes 12). Among these, the phenylpropanoid/shikimate pathway, and more specifically the flavonoid subclass, emerged as the primary short-wavelength-responsive metabolic domain, consistent with findings in cannabis leaves (Kotiranta et al., 2024), *Brassica napus* (Lee et al., 2022), and *Oryza sativa* (Zhang et al., 2024). This selective responsiveness underscores the role of flavonoids as key mediators of plant acclimation to short-wavelength-associated stressors, rather than a broad metabolic response.

A key finding of this work enabled by implementing feature-based molecular networking and MS2LDA substructure discovery, is that UVB selectively increases the abundance of glycosylated flavonoids, while the corresponding aglycone backbones remain largely unchanged. Glycosylation of flavonoids can occur via an oxygen or carbon bond, namely O-and C-glycosylation, respectively, and these post-biosynthetic modifications are key for flavonoids accumulation and auto-toxicity prevention (Behr et al., 2020). Both O-glycosylated flavonols and C-glycosylated flavones were enhanced under UVB treatment. Flavonol glycosides (*e.g.* quercetin-3-O-rutinoside) are stronger antioxidants than flavone glycosides (*e.g.* luteolin-7-O-glucoside), therefore plants prioritize the biosynthesis of flavonols over flavones as quenchers of potential UVB damage (Tattini et al., 2004). Moreover, O-glycosylation is typically associated with vacuolar accumulation of flavonols and fast de-glycosylation, whereas C-glycosylated flavones are more chemically stable and resistant to hydrolysis, suggesting complementary protective functions (Behr et al., 2020; Supplementary Notes 13). The simultaneous accumulation of these structurally and biosynthetically distinct glycosylation types indicates that UVB does not selectively activate a single route, but rather promotes broad flavonoid glycosylation, involving distinct enzyme families with different substrate specificities and cellular localizations as part of a major UV-and light-regulated network of the phenylpropanoid pathway (Le Roy et al., 2016).

The enrichment of glycosylated flavonoids under UVB exposure may reflect a scenario in which cannabis inflorescences increase their capacity to buffer ROS through compartmentalized glycosylated flavonoid pools, which upon oxidative cues are de-glycosylated to rapidly release aglycones capable of quenching ROS, thereby mitigating photon-induced cellular damage. The coordinated increase of both flavonoid subclasses suggests upstream regulation at the transcriptional level of multiple glycosyltransferase pathways, likely via the interaction between UVR8, HY5 and MYB transcription factors, whose binding elements have been localized in glycosyltransferase promoters in rice (Clayton et al., 2018; Zhang et al., 2020). The presence of magenta pigmentation in cannabis glandular trichomes under UVB treatment further supports the notion that flavonoid accumulation contributes to localized photoprotection in reproductive tissues. A yet unannotated anthocyanin-like feature (ID: 2627) (Figure 3D; 4) with a putative glycosylation, was strongly enhanced by UVB, making it a putative candidate responsible for this pigmentation, distinct from the cyanidin-3-rutinoside previously attributed to purple hues in cannabis (Bassolino et al., 2023).

The accumulation of such highly modified flavonoids suggests that UVB exposure not only increases flavonoid abundance but also expands structural complexity, likely enhancing functional diversity and stress robustness. UVB can therefore be used to steer the production of flavonoids in plants, impacting their medicinal and nutritional quality (Contreras-Avilés et al., 2024), as reported in *Fagopyrum esculentum* sprouts (Tian et al., 2024), *Ocimum basilicum* (Skowron et al., 2024), micro-tomato leaves (Lima et al., 2023), and others (Li et al., 2021; Zhu et al., 2023).

Interestingly, the lack of detectable changes in cannabinoid abundances across short-wavelength treatments contrasts with earlier measurements showing that UV radiation broadly enhances cannabis specialized metabolism (Jenkins, 2021; Li et al., 2025; Lydon et al., 1987), reinforcing the idea that cannabinoids and flavonoids fulfil distinct physiological roles in UV-stress adaptation (Agati & Tattini, 2010; Li et al., 2025). Notably, cannflavins were not detected in the chemotype II genotype used, a discussion of the absence is provided in Supplementary Information 14.

Beyond physiological insights, our study highlights how integrated computational metabolomics can connect environmental triggers (*e.g.*, light) to pathway-level classes and post-biosynthetic enzymatic modifications. The combination of FBMN, MS2LDA, SIRIUS/CANOPUS, and deep learning similarity methods (MS2Query and DreaMS) provided the multi-layer evidence needed to interpret treatment effects at the level of flavonoid subclasses and glycosylation types, where exact library matches alone would have resulted in incomplete annotations (Van Der Hooft et al., 2011). Structural assignments should nonetheless be regarded as putative, as flavonoid isomers can remain indistinguishable by MS/MS alone, and mechanistic claims regarding compartmentalization and regulation will benefit from integration with transcriptomic data and targeted MS/MS of diagnostic glycosides.

## Conclusions

In conclusion, our results demonstrate that UVB radiation selectively reshapes the cannabis floral metabolome by promoting the accumulation and structural diversification of flavonoids without any apparent effect on cannabinoids, while UVA does not substantially affect the cannabis floral flavonoid metabolome. The concurrent induction of C- and O-glycosylated flavonoids, supported by substructure-level metabolomic evidence, indicates that UVB-driven acclimation involves coordinated regulation of flavonoid post-biosynthetic modification pathways, advancing our understanding of UV- and light-mediated metabolic plasticity in cannabis. The integrative metabolomics framework developed here can be applied to characterize plant metabolic responses to environmental conditions beyond light, across other plant tissues and natural product classes. Finally, our findings suggest that UVB supplementation in controlled-environment cannabis cultivation may be used to enhance glycosylated flavonoid accumulation, potentially impacting inflorescence medicinal value independently of cannabinoid content, opening new avenues for fine-tuning light spectra in controlled environments to optimize for the presence of specific chemicals in medicinal cannabis.

## Author contributions

W.C.A., L.R.T.O., L.F.M., I.F.K., and E.H. conceived and designed the project. W.C.A. grew the plants, performed the experiments, and collected the data. L.R.T.O. and J.J.J.vdH designed the computational metabolomics workflow. L.R.T.O. built the computational workflow for metabolomics data processing and analysis. W.C.A. and L.R.T.O. contributed equally to data analysis and manuscript preparation. J.J.J.vdH. and I.F.K. supervised the metabolomics component of the project. All authors reviewed and approved the final manuscript.

## Data availability

The authors confirm that the data supporting the findings of this study are available within the article, supporting information, or dedicated repositories. All raw and processed metabolomics data are available in the MASSIVE ID: MSV000100919 including the pre-processed files, metadata and quantification tables. The batch file (mzmine), final annotation table for the flavonoid molecules, GNPS networks (graphML files, job IDs: positive 11072d72767f4e73923333bb0661d5c9 and negative 9a3d6eaf1463432c9fb00c31830031ef), SIRIUS, MS2Query, DreaMS and MS2LDA results, and script used are available at Zenodo (ID: 18715236, https://zenodo.org/records/18715236).

## Acknowledgements

W.C.A. gratefully acknowledges the National Secretariat of Science, Technology, and Innovation of the Republic of Panama (SENACYT, grant no. 270-2020-134). L.R.T.O gratefully recognizes the Marie Skłodowska-Curie grant under the European Union’s Horizon Europe programme MAGiC-MOLFUN (grant no. No. 101072485). We express our gratitude to Gerrit Stunnenberg, David Brink, Dieke Smit, Katina Medina, Marcio León Both, Zacharenia Vlachaki (all Wageningen University and Research) for their technical assistance during the execution cultivation cycle of medicinal cannabis plants under short-wavelengths; Bert Schipper and Shared Research Facilities from Wageningen University and Research for UPLC-MS/MS analysis, and Esteban Charría-Girón for guidance during the metabolomic analysis. Additionally, we acknowledge Marc Juarez in representation of Seoul Semiconductors for providing the UVA and blue LEDs.

## References

1. Agati, G., & Tattini, M. (2010). Multiple functional roles of flavonoids in photoprotection. New Phytologist, 186(4), 786–793. 10.1111/j.1469-8137.2010.03269.x

2. Aliferis, K. A., & Bernard-Perron, D. (2020). Cannabinomics: Application of Metabolomics in Cannabis (Cannabis sativa L.) Research and Development. Frontiers in Plant Science, 11, 554. 10.3389/fpls.2020.00554

3. Bassolino, L., Fulvio, F., Pastore, C., Pasini, F., Gallina Toschi, T., Filippetti, I., & Paris, R. (2023). When Cannabis sativa L. Turns Purple: Biosynthesis and Accumulation of Anthocyanins. Antioxidants, 12(7), 1393. 10.3390/antiox12071393

4. Behr, M., Neutelings, G., El Jaziri, M., & Baucher, M. (2020). You Want it Sweeter: How Glycosylation Affects Plant Response to Oxidative Stress. Frontiers in Plant Science, 11, 571399. 10.3389/fpls.2020.571399

5. Bushuiev, R., Bushuiev, A., Samusevich, R., Brungs, C., Sivic, J., & Pluskal, T. (2025). Self-supervised learning of molecular representations from millions of tandem mass spectra using DreaMS. Nature Biotechnology. 10.1038/s41587-025-02663-3

6. Cerrato, A., Biancolillo, A., Cannazza, G., Cavaliere, C., Citti, C., Laganà, A., Marini, F., Montanari, M., Montone, C. M., Paris, R., Virzì, N., & Capriotti, A. L. (2023). Untargeted cannabinomics reveals the chemical differentiation of industrial hemp based on the cultivar and the geographical field location. Analytica Chimica Acta, 1278, 341716. 10.1016/j.aca.2023.341716

7. Citti, C., Linciano, P., Panseri, S., Vezzalini, F., Forni, F., Vandelli, M. A., & Cannazza, G. (2019). Cannabinoid Profiling of Hemp Seed Oil by Liquid Chromatography Coupled to High-Resolution Mass Spectrometry. Frontiers in Plant Science, 10, 120. 10.3389/fpls.2019.00120

8. Contreras-Avilés, W., Heuvelink, E., Marcelis, L. F. M., & Kappers, I. F. (2024). Ménage à trois: Light, terpenoids, and quality of plants. Trends in Plant Science, 29(5), 572–588. 10.1016/j.tplants.2024.02.007

9. De Jonge, N. F., Louwen, J. J. R., Chekmeneva, E., Camuzeaux, S., Vermeir, F. J., Jansen, R. S., Huber, F., & Van Der Hooft, J. J. J. (2023). MS2Query: Reliable and scalable MS2 mass spectra-based analogue search. Nature Communications, 14(1), 1752. 10.1038/s41467-023-37446-4

10. Desaulniers Brousseau, V., Wu, B.-S., MacPherson, S., Morello, V., & Lefsrud, M. (2021). Cannabinoids and Terpenes: How Production of Photo-Protectants Can Be Manipulated to Enhance Cannabis sativa L. Phytochemistry. Frontiers in Plant Science, 12, 620021. 10.3389/fpls.2021.620021

11. Dührkop, K., Fleischauer, M., Ludwig, M., Aksenov, A. A., Melnik, A. V., Meusel, M., Dorrestein, P. C., Rousu, J., & Böcker, S. (2019). SIRIUS 4: A rapid tool for turning tandem mass spectra into metabolite structure information. Nature Methods, 16(4), 299–302. 10.1038/s41592-019-0344-8

12. Eichholz, I., Rohn, S., Gamm, A., Beesk, N., Herppich, W. B., Kroh, L. W., Ulrichs, C., & Huyskens-Keil, S. (2012). UV-B-mediated flavonoid synthesis in white asparagus (Asparagus officinalis L.). Food Research International, 48(1), 196–201. 10.1016/j.foodres.2012.03.008

13. Ferrer, I. (2020). Analyses of cannabinoids in hemp oils by LC/Q-TOF-MS. In Comprehensive Analytical Chemistry (Vol. 90, pp. 415–452). Elsevier. 10.1016/bs.coac.2020.04.014

14. Fraser, D. P., Sharma, A., Fletcher, T., Budge, S., Moncrieff, C., Dodd, A. N., & Franklin, K. A. (2017). UV-B antagonises shade avoidance and increases levels of the flavonoid quercetin in coriander (Coriandrum sativum). Scientific Reports, 7(1), 17758. 10.1038/s41598-017-18073-8

15. Jenkins, M. W. (2021). &lt;i&gt;Cannabis sativa&lt;/i&gt; L. Response to Narrow Bandwidth UV and the Combination of Blue and Red Light during the Final Stages of Flowering on Leaf Level Gas-Exchange Parameters, Secondary Metabolite Production, and Yield. Agricultural Sciences, 12(12), 1414–1432. 10.4236/as.2021.1212090

16. Jin, D., Dai, K., Xie, Z., & Chen, J. (2020). Secondary Metabolites Profiled in Cannabis Inflorescences, Leaves, Stem Barks, and Roots for Medicinal Purposes. Scientific Reports, 10(1), 3309. 10.1038/s41598-020-60172-6

17. Kim, H. W., Wang, M., Leber, C. A., Nothias, L.-F., Reher, R., Kang, K. B., Van Der Hooft, J. J. J., Dorrestein, P. C., Gerwick, W. H., & Cottrell, G. W. (2021). NPClassifier: A Deep Neural Network-Based Structural Classification Tool for Natural Products. Journal of Natural Products, 84(11), 2795–2807. 10.1021/acs.jnatprod.1c00399

18. Kokalj, D., Zlatić, E., Cigić, B., Kobav, M. B., & Vidrih, R. (2019). Postharvest flavonol and anthocyanin accumulation in three apple cultivars in response to blue-light-emitting diode light. Scientia Horticulturae, 257, 108711. 10.1016/j.scienta.2019.108711

19. Kotiranta, S., Pihlava, J.-M., Kotilainen, T., & Palonen, P. (2024). The morphology, inflorescence yield, and secondary metabolite accumulation in hemp type Cannabis sativa can be influenced by the R:FR ratio or the amount of short wavelength radiation in a spectrum. Industrial Crops and Products, 208, 117772. 10.1016/j.indcrop.2023.117772

20. Le Roy, J., Huss, B., Creach, A., Hawkins, S., & Neutelings, G. (2016). Glycosylation Is a Major Regulator of Phenylpropanoid Availability and Biological Activity in Plants. Frontiers in Plant Science, 7. 10.3389/fpls.2016.00735

21. Li, M., Ouyang, Y., Cai, S., Zhang, Z., Wu, Q., Li, S., Huang, X., Huang, S., Ao, D., You, S., Cai, J., Vasseur, L., Lin, C., & Liu, Y. (2025). A New UV-B Protectant in Plant: Insights Into THCAS and THCA’s Role in UV-B Tolerance. Plant, Cell & Environment, pce.70057. 10.1111/pce.70057

22. Li, T., Yamane, H., & Tao, R. (2021). Preharvest long-term exposure to UV-B radiation promotes fruit ripening and modifies stage-specific anthocyanin metabolism in highbush blueberry. Horticulture Research, 8(1), 67. 10.1038/s41438-021-00503-4

23. Lim, Y. J., Lyu, J. I., Kwon, S.-J., & Eom, S. H. (2021). Effects of UV-A radiation on organ-specific accumulation and gene expression of isoflavones and flavonols in soybean sprout. Food Chemistry, 339, 128080. 10.1016/j.foodchem.2020.128080

24. Lima, I. H. A., Rodrigues, A. A., Resende, E. C., Da Silva, F. B., Farnese, F. D. S., Silva, L. D. J., Rosa, M., Reis, M. N. O., Bessa, L. A., De Oliveira, T. C., Januário, A. H., & Silva, F. G. (2023). Light means power: Harnessing light spectrum and UV-B to enhance photosynthesis and rutin levels in microtomato plants. Frontiers in Plant Science, 14, 1261174. 10.3389/fpls.2023.1261174

25. Liu, Y., Li, S., Shen, M., Guo, F., Li, M., Cai, S., Huang, J., Wu, J., Li, X., Peng, L., Huang, M., Ao, D., Zhu, X., You, S., & Liu, Y. (2025). Integration of transcriptome and metabolome provides insights into metabolites and pathways associated with antiproliferative activity of cannabis flower extracts. Industrial Crops and Products, 223, 120239. 10.1016/j.indcrop.2024.120239

26. Ludwig, M., Nothias, L.-F., Dührkop, K., Koester, I., Fleischauer, M., Hoffmann, M. A., Petras, D., Vargas, F., Morsy, M., Aluwihare, L., Dorrestein, P. C., & Böcker, S. (2020). Database-independent molecular formula annotation using Gibbs sampling through ZODIAC. Nature Machine Intelligence, 2(10), 629–641. 10.1038/s42256-020-00234-6

27. Lydon, J., Teramura, A. H., & Coffman, C. B. (1987). UV-B RADIATION EFFECTS ON PHOTOSYNTHESIS, GROWTH and CANNABINOID PRODUCTION OF TWO Cannabis sativa CHEMOTYPES. Photochemistry and Photobiology, 46(2), 201–206. 10.1111/j.1751-1097.1987.tb04757.x

28. Mao, Y., Luo, J., & Cai, Z. (2025). Biosynthesis and Regulatory Mechanisms of Plant Flavonoids: A Review. Plants, 14(12), 1847. 10.3390/plants14121847

29. Markovitsi, D., Sage, E., Lewis, F. D., & Davies, J. (2013). Interaction of UV radiation with DNA. Photochemical & Photobiological Sciences, 12(8), 1256–1258. 10.1039/c3pp90021e

30. Marti, G., Schnee, S., Andrey, Y., Simoes-Pires, C., Carrupt, P.-A., Wolfender, J.-L., & Gindro, K. (2014). Study of Leaf Metabolome Modifications Induced by UV-C Radiations in Representative Vitis, Cissus and Cannabis Species by LC-MS Based Metabolomics and Antioxidant Assays. Molecules, 19(9), 14004–14021. 10.3390/molecules190914004

31. McCree, K. J. (1981). Photosynthetically Active Radiation. In O. L. Lange, P. S. Nobel, C. B. Osmond, & H. Ziegler (Eds.), Physiological Plant Ecology I: Responses to the Physical Environment (pp. 41–55). Springer. 10.1007/978-3-642-68090-8_3

32. Miao, N., Yun, C., Shi, Y., Gao, Y., Wu, S., Zhang, Z., Han, S., Wang, H., & Wang, W. (2022). Enhancement of flavonoid synthesis and antioxidant activity in Scutellaria baicalensis aerial parts by UV-A radiation. Industrial Crops and Products, 187, 115532. 10.1016/j.indcrop.2022.115532

33. Moreira-Rodríguez, M., Nair, V., Benavides, J., Cisneros-Zevallos, L., & Jacobo-Velázquez, D. (2017). UVA, UVB Light Doses and Harvesting Time Differentially Tailor Glucosinolate and Phenolic Profiles in Broccoli Sprouts. Molecules, 22(7), 1065. 10.3390/molecules22071065

34. Muller, M., & De Villiers, A. (2025). Comprehensive two-dimensional liquid chromatographic analysis of Cannabis phenolics and first evidence of flavoalkaloids in Cannabis. Journal of Chromatography A, 1754, 466023. 10.1016/j.chroma.2025.466023

35. Muthusamy, M., Kim, J. H., Kim, S. H., Kim, J. Y., Heo, J. W., Lee, H., Lee, K.-S., Seo, W. D., Park, S., Kim, J. A., & Lee, S. I. (2020). Changes in Beneficial C-glycosylflavones and Policosanol Content in Wheat and Barley Sprouts Subjected to Differential LED Light Conditions. Plants, 9(11), 1502. 10.3390/plants9111502

36. Myoli, A., Choene, M., Kappo, A. P., Madala, N. E., Van Der Hooft, J. J. J., & Tugizimana, F. (2024a). Charting the Cannabis plant chemical space with computational metabolomics. Metabolomics, 20(3), 62. 10.1007/s11306-024-02125-y

37. Myoli, A., Choene, M., Kappo, A. P., Madala, N. E., Van Der Hooft, J. J. J., & Tugizimana, F. (2024b). Charting the Cannabis plant chemical space with computational metabolomics. Metabolomics, 20(3), 62. 10.1007/s11306-024-02125-y

38. Neugart, S., & Bumke-Vogt, C. (2021). Flavonoid Glycosides in Brassica Species Respond to UV-B Depending on Exposure Time and Adaptation Time. Molecules, 26(2), 494. 10.3390/molecules26020494

39. Neugart, S., & Schreiner, M. (2018). UVB and UVA as eustressors in horticultural and agricultural crops. Scientia Horticulturae, 234, 370–381. 10.1016/j.scienta.2018.02.021

40. Nishigori, C., Yamano, N., Kunisada, M., Nishiaki-Sawada, A., Ohashi, H., & Igarashi, T. (2023). Biological Impact of Shorter Wavelength Ultraviolet RADIATION-C^†^. Photochemistry and Photobiology, 99(2), 335–343. 10.1111/php.13742

41. Nothias, L.-F., Petras, D., Schmid, R., Dührkop, K., Rainer, J., Sarvepalli, A., Protsyuk, I., Ernst, M., Tsugawa, H., Fleischauer, M., Aicheler, F., Aksenov, A. A., Alka, O., Allard, P.-M., Barsch, A., Cachet, X., Caraballo-Rodriguez, A. M., Da Silva, R. R., Dang, T., … Dorrestein, P. C. (2020). Feature-based molecular networking in the GNPS analysis environment. Nature Methods, 17(9), 905–908. 10.1038/s41592-020-0933-6

42. Opálková, M., Navrátil, M., Špunda, V., Blanc, P., & Wald, L. (2018). A database of 10 min average measurements of solar radiation and meteorological variables in Ostrava, Czech Republic. Earth System Science Data, 10(2), 837–846. 10.5194/essd-10-837-2018

43. Palma, C. F. F., Castro-Alves, V., Rosenqvist, E., Ottosen, C., Strid, Å., & Morales, L. O. (2021). Effects of UV radiation on transcript and metabolite accumulation are dependent on monochromatic light background in cucumber. Physiologia Plantarum, 173(3), 750–761. 10.1111/ppl.13551

44. Patil, J. R., Mhatre, K. J., Yadav, K., Yadav, L. S., Srivastava, S., & Nikalje, G. C. (2024). Flavonoids in plant-environment interactions and stress responses. Discover Plants, 1(1), 68. 10.1007/s44372-024-00063-6

45. Rai, N., Morales, L. O., & Aphalo, P. J. (2021). Perception of solar UV radiation by plants: Photoreceptors and mechanisms. Plant Physiology, 186(3), 1382–1396. 10.1093/plphys/kiab162

46. Rai, N., Neugart, S., Yan, Y., Wang, F., Siipola, S. M., Lindfors, A. V., Winkler, J. B., Albert, A., Brosché, M., Lehto, T., Morales, L. O., & Aphalo, P. J. (2019). How do cryptochromes and UVR8 interact in natural and simulated sunlight? Journal of Experimental Botany, 70(18), 4975–4990. 10.1093/jxb/erz236

47. Rea, K. A., Casaretto, J. A., Al-Abdul-Wahid, M. S., Sukumaran, A., Geddes-McAlister, J., Rothstein, S. J., & Akhtar, T. A. (2019). Biosynthesis of cannflavins A and B from Cannabis sativa L. Phytochemistry, 164, 162–171. 10.1016/j.phytochem.2019.05.009

48. Rechner, O., Neugart, S., Schreiner, M., Wu, S., & Poehling, H.-M. (2016). Different Narrow-Band Light Ranges Alter Plant Secondary Metabolism and Plant Defense Response to Aphids. Journal of Chemical Ecology, 42(10), 989–1003. 10.1007/s10886-016-0755-2

49. Shannon, P., Markiel, A., Ozier, O., Baliga, N. S., Wang, J. T., Ramage, D., Amin, N., Schwikowski, B., & Ideker, T. (2003). Cytoscape: A Software Environment for Integrated Models of Biomolecular Interaction Networks. Genome Research, 13(11), 2498–2504. 10.1101/gr.1239303

50. Shomali, A., Das, S., Arif, N., Sarraf, M., Zahra, N., Yadav, V., Aliniaeifard, S., Chauhan, D. K., & Hasanuzzaman, M. (2022). Diverse Physiological Roles of Flavonoids in Plant Environmental Stress Responses and Tolerance. Plants, 11(22), 3158. 10.3390/plants11223158

51. Skowron, E., Trojak, M., & Pacak, I. (2024). Effects of UV-B and UV-C Spectrum Supplementation on the Antioxidant Properties and Photosynthetic Activity of Lettuce Cultivars. International Journal of Molecular Sciences, 25(17), 9298. 10.3390/ijms25179298

52. Sun, X., Kaiser, E., Aphalo, P. J., Marcelis, L. F. M., & Li, T. (2024). Plant responses to UV-A1 radiation are genotype and background irradiance dependent. Environmental and Experimental Botany, 219, 105621. 10.1016/j.envexpbot.2023.105621

53. Sun, X., Kaiser, E., Zhang, Y., Marcelis, L. F. M., & Li, T. (2025). Quantifying the Photosynthetic Quantum Yield of Ultraviolet-A1 Radiation. Plant, Cell & Environment, 48(1), 109–121. 10.1111/pce.15145

54. Tang, Q., Xu, Y., Gao, F., Xu, Y., Cheng, C., Deng, C., Chen, J., Yuan, X., Zhang, X., & Su, J. (2023). Transcriptomic and metabolomic analyses reveal the differential accumulation of phenylpropanoids and terpenoids in hemp autotetraploid and its diploid progenitor. BMC Plant Biology, 23(1), 616. 10.1186/s12870-023-04630-z

55. Tattini, M., Galardi, C., Pinelli, P., Massai, R., Remorini, D., & Agati, G. (2004). Differential accumulation of flavonoids and hydroxycinnamates in leaves of *Ligustrum vulgare* under excess light and drought stress. New Phytologist, 163(3), 547–561. 10.1111/j.1469-8137.2004.01126.x

56. Tian, X., Hu, M., Yang, J., Yin, Y., & Fang, W. (2024). Ultraviolet-B Radiation Stimulates Flavonoid Biosynthesis and Antioxidant Systems in Buckwheat Sprouts. Foods, 13(22), 3650. 10.3390/foods13223650

57. Torres Ortega, L. R., Dietrich, J., Wandy, J., Mol, H., & Van Der Hooft, J. J. J. (2025). Large-scale discovery and annotation of hidden substructure patterns in mass spectrometry profiles. Biochemistry. 10.1101/2025.06.19.659491

58. Vásquez-Ocmín, P. G., Marti, G., Bonhomme, M., Mathis, F., Fournier, S., Bertani, S., & Maciuk, A. (2021). Cannabinoids vs. whole metabolome: Relevance of cannabinomics in analyzing Cannabis varieties. Analytica Chimica Acta, 1184, 339020. 10.1016/j.aca.2021.339020

59. Zhang, X., Ding, X., Ji, Y., Wang, S., Chen, Y., Luo, J., Shen, Y., & Peng, L. (2018). Measurement of metabolite variations and analysis of related gene expression in Chinese liquorice (Glycyrrhiza uralensis) plants under UV-B irradiation. Scientific Reports, 8(1), 6144. 10.1038/s41598-018-24284-4

60. Zhao, B., Wang, L., Pang, S., Jia, Z., Wang, L., Li, W., & Jin, B. (2020). UV-B promotes flavonoid synthesis in Ginkgo biloba leaves. Industrial Crops and Products, 151, 112483. 10.1016/j.indcrop.2020.112483

61. Zhu, X., Trouth, F., & Yang, T. (2023). Preharvest UV-B Treatment Improves Strawberry Quality and Extends Shelf Life. Horticulturae, 9(2), 211. 10.3390/horticulturae9020211

