## Supplementary Information for "Characterizing the effect of short wavelengths on the floral flavonoid metabolome of medicinal cannabis using a comparative computational metabolomics workflow"

### Introduction

**Supplementary Notes 1:** The electromagnetic spectrum of solar radiation reaching the terrestrial surfaces ranges from 100 to 1x$10^6$ nm, encompassing the photosynthetically active radiation (PAR, 400-700 nm) (McCree, 1981) and ultraviolet radiation (UV, 100-400 nm), among others (Björn, 2015). Outside of the PAR spectrum, UVA waveband is often divided into short (315-350 nm) and long wavelength (350-400 nm) as these are perceived by UVR8 and CRYs, respectively (Rai et al., 2021).

**Supplementary Notes 2:** Humans depend on the consumption of fruits and vegetables for the intake of flavonoids that improve vascular (Khani & Spencer, 2025) and cognitive functions (Cheng et al., 2022), as well as prevent diseases associated to metabolic syndrome (Gouveia et al., 2022).

**Supplementary Notes 3:** Flavonoids are mostly produced in plants, but also by bacteria and fungi, and their structural subunit consists of a diphenylpropane C6-C3-C6 core of aromatic rings (A- and B-) linked to a heterocyclic C-ring (Davies et al., 2024).

**Supplementary Notes 4:** Glycosylation is the enzymatic conjugation of an aglycone to a sugar moiety (glucose, galactose, rhamnose, xylose, arabinose) catalysed by the action of glycosyltransferases (Gachon et al., 2005; Pandey & Sohng, 2016). A glycosylation makes flavonoid vacuolar sequestration, transport, stabilization, and synthesis feasible (Gharabli et al., 2023).

**Supplementary Notes 5:** Computational metabolomic studies can use targeted and untargeted approaches, where the latter allows to detect considerably greater numbers of unknown compounds that allows for a more robust portrait of the metabolome fingerprint of specific genotypes, developmental stages, tissues/structures, and environmental conditions (Chen et al., 2020). For instance, to systematically investigate how different light conditions reshape the floral flavonoid metabolome, untargeted high-resolution ultra-high performance liquid chromatography coupled to tandem mass spectrometry (UPLC–MS/MS) was used, typically yielding thousands of chromatographic features (*m/z*–retention time pairs) with associated fragmentation spectra for comparative analyses across treatments (Hakeem et al., 2025; Kalinski et al., 2026). However, translating these signals into confident chemical identities remains a key bottleneck, particularly for flavonoids and their glycosylated patterns, which often occur as positional isomers producing highly similar MS/MS patterns (Li et al., 2024). Moreover, exact spectral-library matching frequently provides limited coverage because many experimental spectra have no direct reference, motivating approaches that expand annotation beyond exact matches (Bittremieux et al., 2022).

To overcome this annotation bottleneck, we created and applied an integrated computational workflow that links differential abundance patterns to molecular families, substructures, and candidate annotations to study the effects of short wavelengths on cannabis flower metabolomes. Specifically, molecular networking (MN) organizes MS/MS spectra into molecular families to prioritize treatment-responsive clusters and support annotation propagation across related metabolites. Subsequently, MS2LDA 2.0 identifies recurring fragmentation motifs (Mass2Motifs) to explore substructures and moieties that differentiate light conditions (Torres Ortega et al., 2025) while SIRIUS supports molecular-formula inference, chemical class assignments, and structure-candidate generation (Dührkop et al., 2019). Finally, MS2Query (De Jonge et al., 2023) and DreaMS (Bushuiev et al., 2025) are machine learning and deep learning similarity based-approaches that were used to retrieve analogs if a reliable spectral match from library was not found.

### Materials and methods

**Supplementary Information 6:**

Glandular trichome imaging: Inflorescence bracts were collected from within 5 cm of the apical section of the top inflorescences and examined for their glandular trichomes. Photographs were made under a stereomicroscope (ZEISS Stemi SV 11, Carl Zeiss, Germany) with a zoom range of 0.6× to 6.6× using dark-field illumination to compare glandular trichome coloration across short-wavelength treatments.

**Supplementary Information 7:**

Ultra-high-performance liquid chromatography mass spectrometry (UHPLC-MS/MS) based metabolomic analysis: For each sample, 10 µL was injected into a UHPLC with Exploris120 Orbitrap system (Thermo Scientific) and vanquish photodiode array detector (220-600 nm). The Orbitrap Exploris120 mass spectrometer was equipped with an electrospray ionization (ESI) source. Chromatographic separation was performed on a reversed-phase column (Acquity UPLC BEH C18, 1.7 µm, 2.1 × 150 mm; Waters) at 40°C with a mobile phase composed of degassed eluent A [ultra-pure water: formic acid (1000:1, v/v)] and eluent B [acetonitrile: formic acid (1000:1, v/v)] at a flow rate of 0.4 mL $min^{-1}$. A linear gradient of 5 to 75% (v/v) eluent B was applied over 22 minutes, followed by 8 minutes of washing and equilibration. UV absorbance was monitored using a Vanquish photodiode array detector (220–600 nm). Fourier transform mass spectrometry (FTMS) full scans (m/z 90.00–1350.00) were recorded at a resolution of 60,000 full width at half maximum (FWHM). ESI was performed in alternating positive and negative modes with spray voltages of +3 kV (ESI+) and −2.5 kV (ESI−).

**Supplementary Information 8:**

Data preprocessing and metabolite feature finding: The mzML files were pre-processed using mzmine 4.7.8 with the mzwizard workflow (batch file available at Zenodo, see Data Availability). Chromatograms were cropped between 0.5 to 60 minutes for retention time. The maximum number of peaks per chromatogram was set to 15, with a minimum of 4 consecutive scans per peak. The minimum feature height was set to $5.04 \times 10^6$ and noise thresholds were MS1 = 5.0 and MS2 = 2.5. Metabolic features with signal intensities less than threefold above those detected in the blank were excluded.

**Supplementary Information 9:**

Metabolite and substructure *in silico* annotation: The settings used for SIRIUS were the default: mass accuracy of 5 ppm, adding the main fallback adducts: [M + H]+, [M + Na]+ and [M + K]+. The databases selected were the plant related types (such as GNPS, Plantcyc, LOTUS, Biocyc). We used SIRIUS molecular formula identification with the '*de novo* + bottom-up' strategy (*m/z* threshold 400): bottom-up candidate generation was applied to all features, and *de novo* formula enumeration was additionally performed only for precursors *m/z* < 400. For DreaMS, we obtained the most similar reference spectra above a cosine similarity threshold of 0.75 (via the DreaMS web interface).

For the MS2LDA run the spectra were filtered to include a minimum of five fragments per spectra, and a minimum normalized intensity of 0.03. For the MS2LDA settings, the number of Mass2Motifs was set to 150 with 10000 iterations and fragments and losses per Mass2Motifs of 50. For the substructure Annotation Guidance, five reference molecules were retrieved with scores above 0.90 of cosine similarity.

**Supplementary Information 10:**

Feature-based molecular networking: for the feature-based molecular networking using GNPS2 (Nothias et al., 2020), the following settings were used: fragment tolerance = 0.5, precursor mass tolerance = 0.02, cosine similarity threshold = 0.7, minimum matched peaks = 6, and library TopK = 2. For molecular networking, edges were created using a cosine similarity cutoff of 0.7 with a minimum of six matched peaks, TopK = 10, and a maximum component size of 100. Altogether, the outputs from SIRIUS (formula/ZODIAC/CANOPUS), MS2Query, DreaMS, and MS2LDA were exported as feature-level tables and integrated with the network by importing them into Cytoscape as node attribute tables and joining them to network nodes using the shared feature identifier. This integration provided additional annotations for network nodes without direct spectral library matches.

**Supplementary Information 11:**

Chemical classification and chemometric analysis: datasets were pre-processed in MetaboAnalyst 13.0 (Pang et al., 2021) as follows: missing values were estimated using left-censored data imputation, features were normalized by sample median, $log_{10}$ transformed, and Pareto scaled prior to multivariate analysis.

Partial least squares discriminant analysis (PLS-DA) was performed on each dataset and validated using five-fold cross-validation. Model performance was assessed by $R^2$, indicating the proportion of variance explained by the model, and $Q^2$, defined as 1 minus the ratio of the predictive residual sum of squares to the total sum of squares of the response variable, indicating predictive performance. Variable importance in projection (VIP) scores were calculated from the PLS-DA model; features with VIP ≥ 1.5 were considered significant contributors to group separation, and features with VIP ≥ 2.5 were considered the strongest drivers.

Hierarchical clustering heatmaps were generated using Ward linkage distance and Z-scaling of feature abundances ranging from -1 to 1. Features displayed in heatmaps were filtered by VIP ≥ 0.5 (*phenylpropanoid* dataset) or VIP ≥ 1.5 (*global* dataset) depending on the analysis.

Mean separation of relative abundances across treatments was performed using a one-way ANOVA followed by Fisher's protected least significant difference (LSD) post-hoc test at $\alpha = 0.05$.

Treatment effect magnitude was assessed by volcano plot analysis using pairwise t-tests comparing each short-wavelength treatment to the reference. A threshold of $-\log_{10}$(p-value) ≥ 1.3 (corresponding to $p < 0.05$) was applied for statistical significance, and a $\log_2$-fold change (FC) threshold of ≥ 2 or ≤ −2 was used to define biologically meaningful magnitude of change. Positive and negative $\log_2$FC values indicate upregulation and downregulation of relative abundances, respectively.

**Supplementary Notes 12:** Blue light upregulates the metabolism of fatty acids in Oryza sativa (Zhang et al., 2024) while downregulating terpenoids in Salvia miltiorrhiza (Chen et al., 2018). Previous studies have shown that UVB enriches the metabolism of amino acids and carbohydrates in *Rhododendron chrysanthum* (Sun et al., 2022), terpenoids in *Artemisia annua* (Rai et al., 2011), and alkaloids in Catharanthus roseus (Zhu et al., 2015); while UVA also upregulates the metabolism of amino acids in *Brassica alboglabra* (He et al., 2021) and terpenoids in *Prunus persica* (Zhang et al., 2025).

**Supplementary Notes 13:** In *Spirodela polyrhiza,* the glycosylated flavone luteolin 8-C-glucoside is mostly accumulated in the vacuole and chloroplast of mesophyll cells conferring copper-sulphate stress tolerance; whereas luteolin 7-O-glucoside accumulates in the vacuole of epidermal cells conferring tolerance to copper-sulphate and UV stress (Böttner et al., 2021).

**Supplementary Notes 14:** Based on our analyses, we could not detect prenylated flavones (also known as cannflavins) in the chemotype II (THC:CBD 1:1) cannabis genotype used. Cannflavin levels are mainly determined by the genetics (genotype) of the plant along with radiation and low air temperatures (6-16 °C) at 1200 m above sea level (Calzolari et al., 2017; Giupponi et al., 2020). Accumulation of cannflavins has been reported in chemotypes I (high THC) (Radwan et al., 2008), as well in chemotypes III (low THC) (Cerrato et al., 2023; Giupponi et al., 2020; Kotiranta et al., 2024). Nevertheless, our evidence is not sufficient to assert that a chemotype II does not produce cannflavins, nor can we further discuss whether the short-wavelength treatments had an influence on their abundances.

### Figures

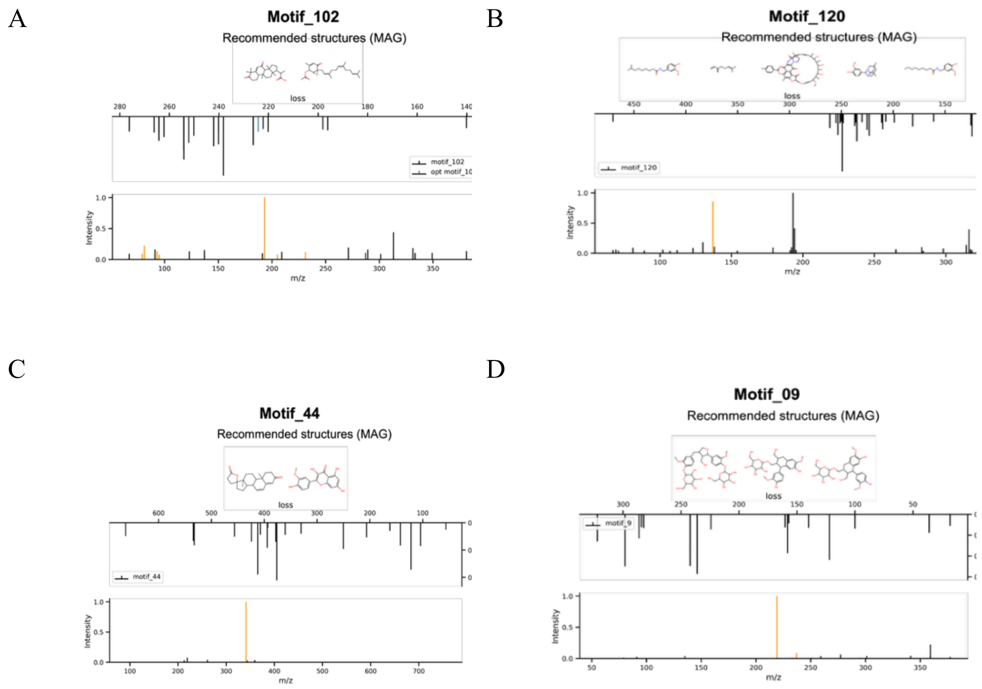

**Supplementary Figure 1 |** (A) motif_102, (B) motif_120, (C) motif_44, and (D) motif_9, corresponding to the cannabinoids CBD, CGB, CDBA and CBGA, respectively.

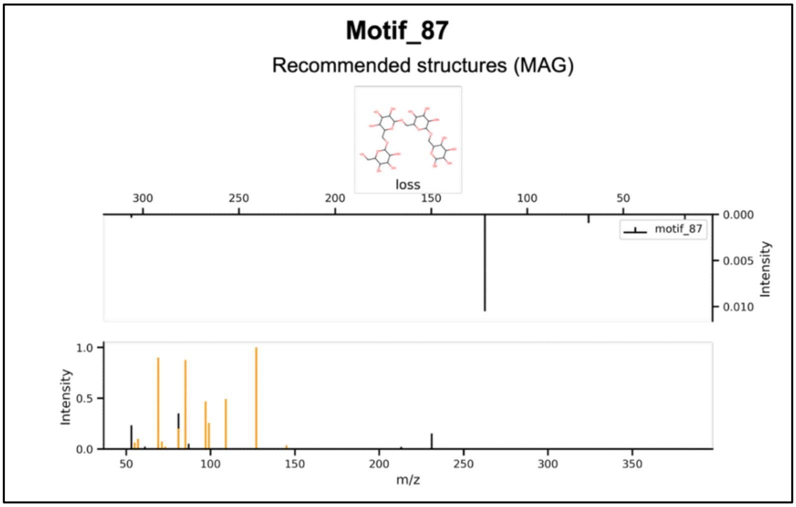

**Supplementary Figure 2 |** Motif_87. On top, the recommended structure given by MAG. The spectrum above refers to the losses contained in the motif. Below, a spectrum with the fragments associated to the motif, in orange the optimized fragments (those existing in the recommended structure and are part of the motif) corresponding to the substructure drawn.

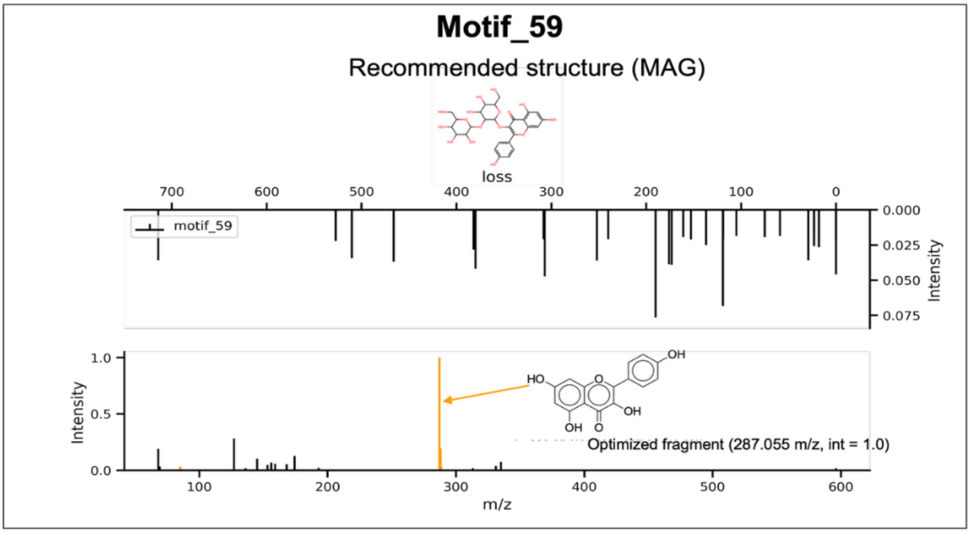

**Supplementary Figure 3 |** Motif_59. On top, the recommended structure given by MAG. The spectrum above refers to the losses contained in the motif. Below, a spectrum with the fragments associated to the motif, in orange the optimized fragments (those existing in the recommended structure and are part of the motif) corresponding to the substructure drawn.

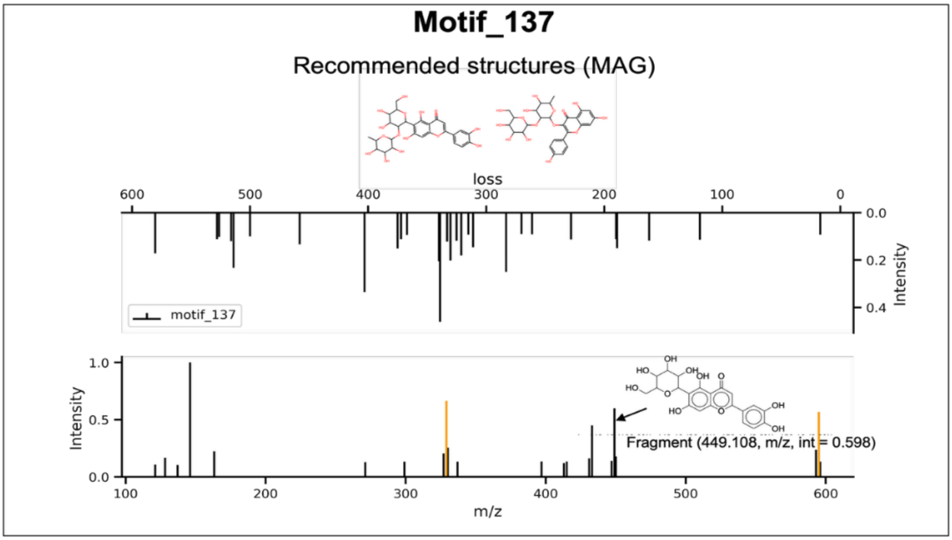

**Supplementary Figure 4 |** (A) motif_137. On top the recommended structures given by MAG. The spectrum above refers to the losses contained in the motif. Below, a spectrum with the fragments associated to the motif, in orange the optimized fragments (those existing in the recommended structure and are part of the motif) corresponding to the substructure drawn.

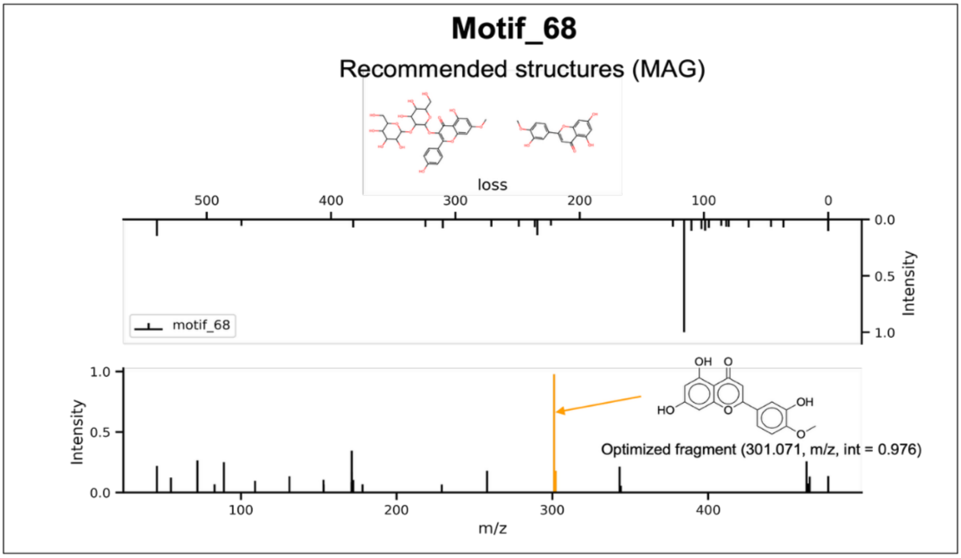

**Supplementary Figure 5 |** Motif_68. On top, the recommended structures given by MAG. The spectrum above refers to the losses contained in the motif. Below, a spectrum with the fragments associated to the motif, in orange the optimized fragments (those existing in the recommended structure and are part of the motif) corresponding to the substructure drawn.

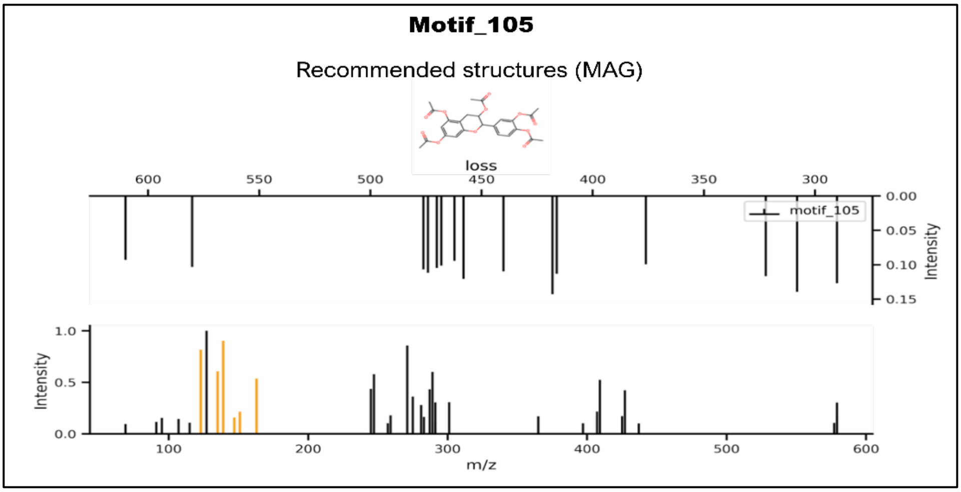

**Supplementary Figure 6 |** Motif_105. On top, the recommended structure given by MAG. The spectrum above refers to the losses contained in the motif. Below, a spectrum with the fragments associated to the motif, in orange the optimized fragments (those existing in the recommended structure and are part of the motif) corresponding to the substructure drawn

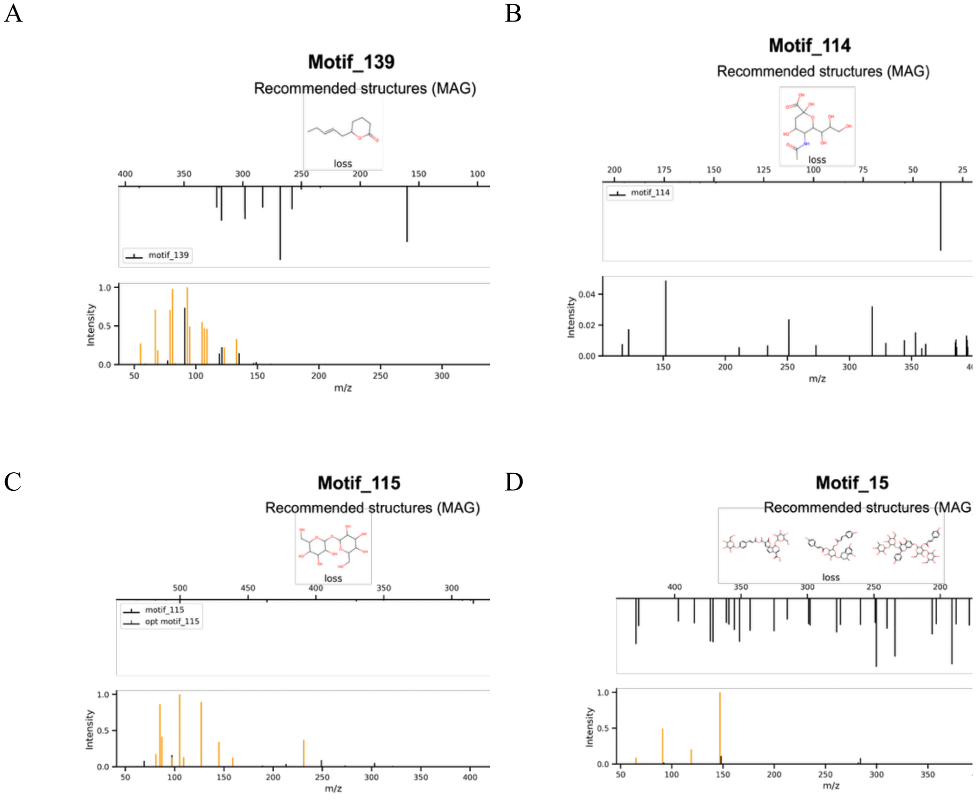

**Supplementary Figure 7 |** (A) motif_139, (B) motif_114, (C) motif_115, and (D) motif_15, associated to glycosylation moieties.

**Supplementary Table 1 |** All putatively annotated flavonoids (level 2 in positive ion mode) in *Cannabis sativa* L. inflorescences.

| ID | *m/z* | Rt (min) | Putative Annotation | MS2Query Score | Motif | MS2Q Class | SIRIUS Class | DreaMS Annotation | DreaMS Similarity | Confidence |
|---|---|---|---|---|---|---|---|---|---|---|
| **426** | 301.070 | 20.88 | Diosmetin | 0.9244 | 68 | Flavones | Flavones | Diosmetin | 0.7788 | High |
| **373** | 287.054 | 18.00 | Luteolin | 0.9343 | 59 | Flavonols | Flavones | Peonidin-3,5-O-di-eta-glucoside | 0.7771 | High |
| **307** | 449.108 | 15.01 | Luteolin-4'-O-glucoside | 0.9334 | 59 | Flavones | Flavones | Luteolin | 0.7660 | Medium |
| **304** | 464.126 | 14.81 | Peonidin-3-O-glucoside | 0.8634 | 68 | — | Purine nucleosides | Diosmetin -7-Orutinoside | — | Medium |
| **291** | 609.145 | 14.32 | Diosmetin-7-O-neohesperidoside | 0.8634 | 68 | Flavones | Flavones | Flavone glycoside | — | Medium |
| **288** | 593.186 | 14.18 | Margaritene | 0.9591 | 137 | Flavones | Flavones | Kaempferide | — | Medium |
| **278** | 287.055 | 13.88 | Kaempferol | 0.9343 | 59 | Flavonols | Flavonols | Isorhamnetin-3-O-glucoside | — | High |
| **269** | 449.107 | 13.79 | Kaempferol-3-O-glucoside | 0.9703 | 59 | Flavonols | Flavonols | Poly-Glycosylated flavonol | — | High |
| **266** | 625.176 | 13.40 | Isorhamnetin-3-O-rutinoside | 0.9614 | 68 | Flavonols | Flavonols | Luteolin-4'-O-beta-D-glucopyranoside | 0.8538 | High |
| **260** | 595.165 | 13.17 | Kaempferol-3-O-rutinoside | 0.9610 | 59 | Flavonols | Flavonols | Isorhamnetin-3-O-glucoside | — | Medium |
| **256** | 463.123 | 12.96 | Hispidulin-4′-O-glucoside | 0.9362 | 68 | Flavones | Flavones | Peonidin-3-O-beta-D-glucopyranoside | — | Medium |
| **253** | 449.107 | 12.75 | Luteolin-7-glucuronide | 0.9536 | 59 | Flavones | Flavones | Diosmin-like flavone | 0.8530 | High |
| **243** | 433.112 | 12.27 | Vitexin | 0.9700 | 137 | Flavones | Flavones | Diosmetin-7-O-neohes | 0.7916 | High |

| | | | | | | | | | | |
|---|---|---|---|---|---|---|---|---|---|---|
| | | | | | | | | peridosi de | | |
| **235** | 579.170 | 11.84 | Vitexin-2 -O-rhamnoside | 0.9600 | 137 | Flavones | Flavones | Margaritene | 0.8728 | High |
| **226** | 449.107 | 11.29 | Orientin | 0.9700 | 137 | Flavones | Flavones | Kaempferol | 0.7677 | Medium |
| **145** | 596.169 | 7.37 | Nicotiflorin | 0.8634 | 59 | Flavonols | — | Vitexin 2' '-O-glucoside | 0.7660 | Low |

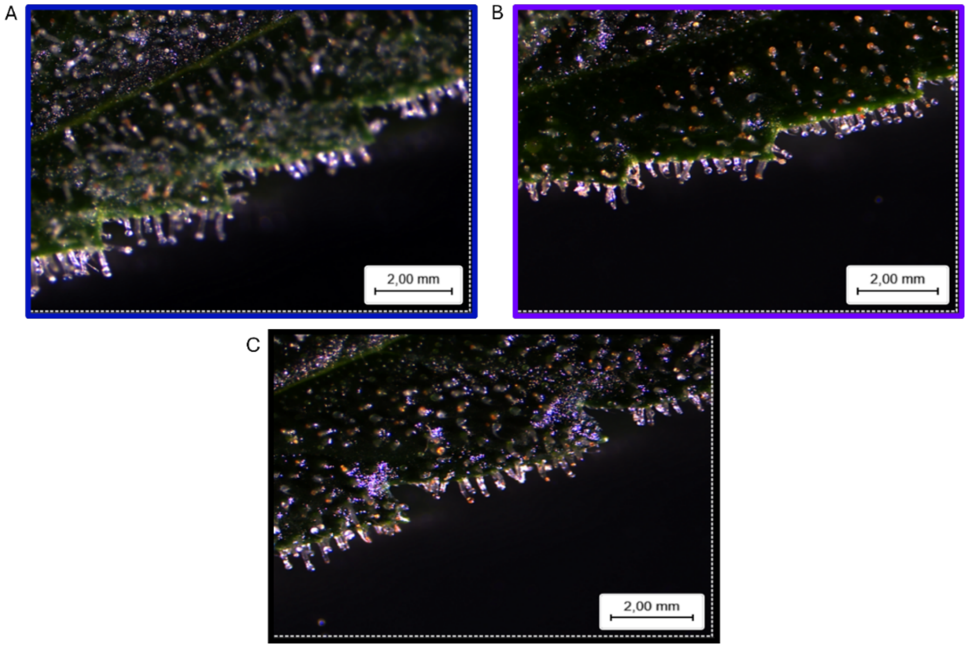

**Supplementary Figure 8 |** Glandular trichomes in inflorescence bracts from the sixth week of the short-day phase of *Cannabis sativa* L under (A) blue, (B) UVA (C) reference treatments.

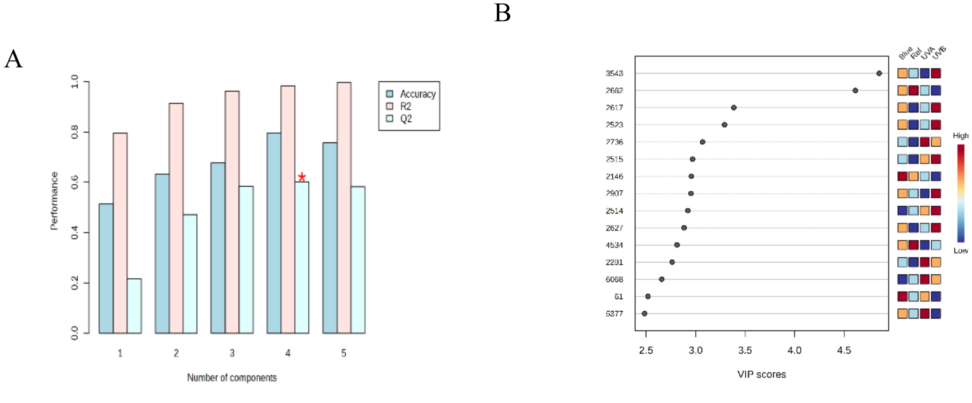

**Supplementary Figure 9 |** (A) Model performance statistics ($R^2$, $Q^2$, and accuracy) obtained by 5-fold cross-validation, and (B) Features with variables in projection (VIP) scores ≥ 2.5 corresponding to the PLS-DA of the '*phenylpropanoid/shikimate'* dataset. A red asterisk (*) indicates that the predictive performance ($Q^2$) of the four-component PLS-DA model is significantly greater than that of permuted models ($p < 0.05$), as determined by permutation testing.

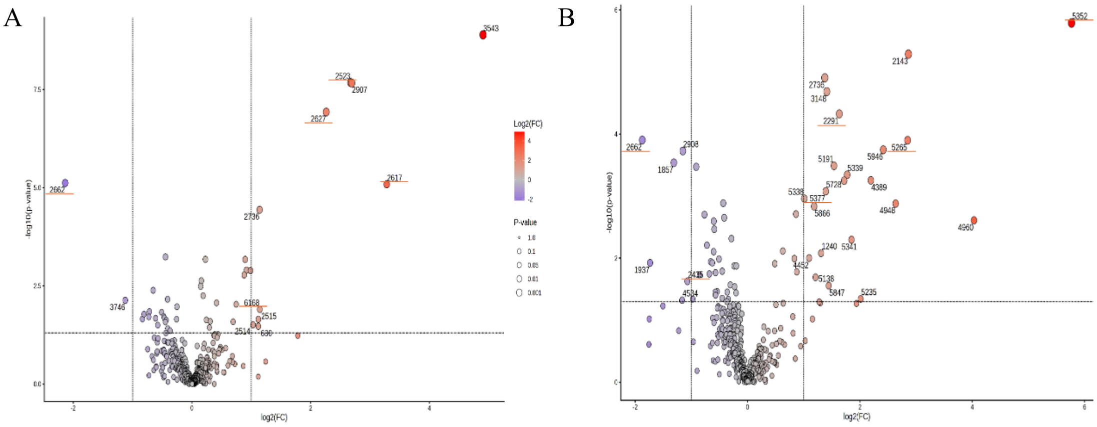

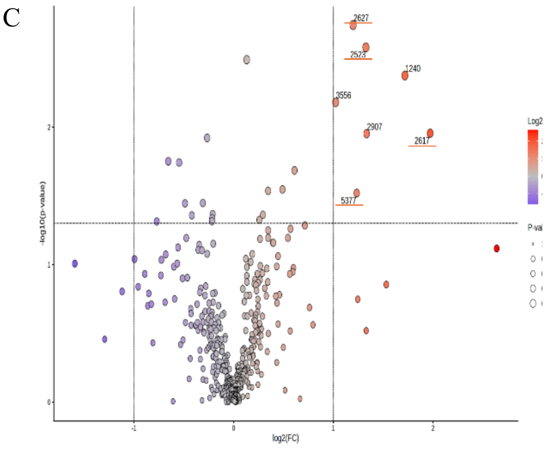

**Supplementary Figure 10 |** Volcano plots displaying relative abundance changes in response to short-wavelengths. (A) UVB vs reference, (B) UVA vs reference, and (C) blue vs reference. X-axis shows $Log_2$ fold-change, positive values indicate higher relative abundance in light treatment. The y-axis shows $-log_{10}$(p-value), higher points indicate stronger statistical significance. Dotted lines indicate fold-change (vertical) and *p*-value (horizontal) cutoffs threshold of 0.05 and 2, respectively. Color scale indicates direction of change (blue = downregulated; red = upregulated), and point size corresponds to the *p*-value. Labeled points (ID number) correspond to features meeting both significance criteria. An orange underscore indicate those features which are flavonoids.

### References

Bittremieux, W., Wang, M., & Dorrestein, P. C. (2022). The critical role that spectral libraries play in capturing the metabolomics community knowledge. *Metabolomics*, *18*(12), 94. https://doi.org/10.1007/s11306-022-01947-y

Björn, L. O. (2015). Ultraviolet-A, B, and C. *UV4Plants Bulletin*, *2015*(1), 17–18. https://doi.org/10.19232/uv4pb.2015.1.12

Böttner, L., Grabe, V., Gablenz, S., Böhme, N., Appenroth, K. J., Gershenzon, J., & Huber, M. (2021). Differential localization of flavonoid glucosides in an aquatic plant implicates different functions under abiotic stress. *Plant, Cell & Environment*, *44*(3), 900–914. https://doi.org/10.1111/pce.13974

Bushuiev, R., Bushuiev, A., Samusevich, R., Brungs, C., Sivic, J., & Pluskal, T. (2025). Self-supervised learning of molecular representations from millions of tandem mass spectra using DreaMS. *Nature Biotechnology*. https://doi.org/10.1038/s41587-025-02663-3

Calzolari, D., Magagnini, G., Lucini, L., Grassi, G., Appendino, G. B., & Amaducci, S. (2017). High added-value compounds from Cannabis threshing residues. *Industrial Crops and Products*, *108*, 558–563. https://doi.org/10.1016/j.indcrop.2017.06.063

Cerrato, A., Biancolillo, A., Cannazza, G., Cavaliere, C., Citti, C., Laganà, A., Marini, F., Montanari, M., Montone, C. M., Paris, R., Virzì, N., & Capriotti, A. L. (2023). Untargeted cannabinomics reveals the chemical differentiation of industrial hemp based on the cultivar and the geographical field location. *Analytica Chimica Acta*, *1278*, 341716. https://doi.org/10.1016/j.aca.2023.341716

Chen, L., Zhong, F., & Zhu, J. (2020). Bridging Targeted and Untargeted Mass Spectrometry-Based Metabolomics via Hybrid Approaches. *Metabolites*, *10*(9), 348. https://doi.org/10.3390/metabo10090348

Cheng, N., Bell, L., Lamport, D. J., & Williams, C. M. (2022). Dietary Flavonoids and Human Cognition: A Meta-Analysis. *Molecular Nutrition & Food Research*, *66*(21), 2100976. https://doi.org/10.1002/mnfr.202100976

Davies, K. M., Andre, C. M., Kulshrestha, S., Zhou, Y., Schwinn, K. E., Albert, N. W., Chagné, D., Van Klink, J. W., Landi, M., & Bowman, J. L. (2024). The evolution of flavonoid biosynthesis. *Philosophical Transactions of the Royal Society B: Biological Sciences*, *379*(1914), 20230361. https://doi.org/10.1098/rstb.2023.0361

De Jonge, N. F., Louwen, J. J. R., Chekmeneva, E., Camuzeaux, S., Vermeir, F. J., Jansen, R. S., Huber, F., & Van Der Hooft, J. J. J. (2023). MS2Query: Reliable and scalable MS2 mass spectra-based analogue search. *Nature Communications*, *14*(1), 1752. https://doi.org/10.1038/s41467-023-37446-4

Dührkop, K., Fleischauer, M., Ludwig, M., Aksenov, A. A., Melnik, A. V., Meusel, M., Dorrestein, P. C., Rousu, J., & Böcker, S. (2019). SIRIUS 4: A rapid tool for turning tandem mass spectra into metabolite structure information. *Nature Methods*, *16*(4), 299–302. https://doi.org/10.1038/s41592-019-0344-8

Gachon, C. M. M., Langlois-Meurinne, M., & Saindrenan, P. (2005). Plant secondary metabolism glycosyltransferases: The emerging functional analysis. *Trends in Plant Science*, *10*(11), 542–549. https://doi.org/10.1016/j.tplants.2005.09.007

Gharabli, H., Della Gala, V., & Welner, D. H. (2023). The function of UDP-glycosyltransferases in plants and their possible use in crop protection. *Biotechnology Advances*, *67*, 108182. https://doi.org/10.1016/j.biotechadv.2023.108182

Giupponi, L., Leoni, V., Pavlovic, R., & Giorgi, A. (2020). Influence of Altitude on Phytochemical Composition of Hemp Inflorescence: A Metabolomic Approach. *Molecules*, *25*(6), 1381. https://doi.org/10.3390/molecules25061381

Gouveia, H. J. C. B., Urquiza-Martínez, M. V., Manhães-de-Castro, R., Costa-de-Santana, B. J. R., Villarreal, J. P., Mercado-Camargo, R., Torner, L., De Souza Aquino, J., Toscano, A. E., & Guzmán-Quevedo, O. (2022). Effects of the Treatment with Flavonoids on Metabolic Syndrome Components in Humans: A Systematic Review Focusing on Mechanisms of Action. *International Journal of Molecular Sciences*, *23*(15), 8344. https://doi.org/10.3390/ijms23158344

Hakeem, M. K., Rajendaran, T., Eldin Saeed, E., Mishra, A. K., Hazzouri, K. M., Shah, I., & Amiri, K. M. A. (2025). Comparative LC-MS/MS-based profiling of phytohormones: A unified analytical approach across diverse plant matrices. *Frontiers in Plant Science*, *16*, 1670979. https://doi.org/10.3389/fpls.2025.1670979

Kalinski, J.-C., Ruiz Brandão da Costa, B., Schramm, T., Buckett, L. R., Carlson, L. T., Coffey, N. R., Damiani, T., Dechent, E., Abiead, Y. E., Heuckeroth, S., Jennings, E. K., Kaesler, J., Stock, N. L., Orme, A. M., Torres, R. R., Trojahn, S., Whelton, H. L., Yan, Y., Aron, A. T., … Petras, D. (2026). Comparability of Liquid Chromatography Tandem Mass Spectrometry Analysis of Dissolved Organic Matter across Laboratories. *Environmental Science & Technology*. https://doi.org/10.1021/acs.est.5c12691

Khani, S., & Spencer, J. (2025). A systematic review of the mechanisms of cardiovascular disease reduction by dietary flavonoids: The impact of anthocyanins on flow-mediated dilation and blood rheology. *BMC Nutrition*, *11*(1), 195. https://doi.org/10.1186/s40795-025-01086-2

Kotiranta, S., Pihlava, J.-M., Kotilainen, T., & Palonen, P. (2024). The morphology, inflorescence yield, and secondary metabolite accumulation in hemp type Cannabis sativa can be influenced by the R:FR ratio or the amount of short wavelength radiation in a spectrum. *Industrial Crops and Products*, *208*, 117772. https://doi.org/10.1016/j.indcrop.2023.117772

Li, W., Zhang, X., Wang, S., Gao, X., & Zhang, X. (2024). Research Progress on Extraction and Detection Technologies of Flavonoid Compounds in Foods. *Foods*, *13*(4), 628. https://doi.org/10.3390/foods13040628

McCree, K. J. (1981). Photosynthetically Active Radiation. In O. L. Lange, P. S. Nobel, C. B. Osmond, & H. Ziegler (Eds.), *Physiological Plant Ecology I: Responses to the Physical Environment* (pp. 41–55). Springer. https://doi.org/10.1007/978-3-642-68090-8_3

Nothias, L.-F., Petras, D., Schmid, R., Dührkop, K., Rainer, J., Sarvepalli, A., Protsyuk, I., Ernst, M., Tsugawa, H., Fleischauer, M., Aicheler, F., Aksenov, A. A., Alka, O., Allard, P.-M., Barsch, A., Cachet, X., Caraballo-Rodriguez, A. M., Da Silva, R. R., Dang, T., … Dorrestein, P. C. (2020). Feature-based molecular networking in the GNPS analysis environment. *Nature Methods*, *17*(9), 905–908. https://doi.org/10.1038/s41592-020-0933-6

Pandey, R. P., & Sohng, J. K. (2016). Glycosyltransferase-Mediated Exchange of Rare Microbial Sugars with Natural Products. *Frontiers in Microbiology*, *7*. https://doi.org/10.3389/fmicb.2016.01849

Radwan, M. M., ElSohly, M. A., Slade, D., Ahmed, S. A., Wilson, L., El-Alfy, A. T., Khan, I. A., & Ross, S. A. (2008). Non-cannabinoid constituents from a high potency Cannabis sativa variety. *Phytochemistry*, *69*(14), 2627–2633. https://doi.org/10.1016/j.phytochem.2008.07.010

Rai, N., Morales, L. O., & Aphalo, P. J. (2021). Perception of solar UV radiation by plants: Photoreceptors and mechanisms. *Plant Physiology*, *186*(3), 1382–1396. https://doi.org/10.1093/plphys/kiab162

Torres Ortega, L. R., Dietrich, J., Wandy, J., Mol, H., & Van Der Hooft, J. J. J. (2025). *Large-scale discovery and annotation of hidden substructure patterns in mass spectrometry profiles*. Biochemistry. https://doi.org/10.1101/2025.06.19.659491
